# Testing for a role of postzygotic incompatibilities in rapidly speciated Lake Victoria cichlids

**DOI:** 10.1101/2023.09.26.559612

**Authors:** Anna F. Feller, Catherine L. Peichel, Ole Seehausen

## Abstract

Intrinsic postzygotic hybrid incompatibilities are usually due to negative epistatic interactions between alleles from different parental genomes. While such incompatibilities are thought to be uncommon in speciation with gene flow, they may be important if such speciation results from a hybrid population. Here we aimed to test this idea in the endemic cichlid fishes of Lake Victoria. Hundreds of species have evolved within the lake in <15k years from a hybrid progenitor. While the importance of prezygotic barriers to gene flow is well established in this system, the possible relevance of postzygotic genetic incompatibilities is unknown. We inferred the presence of negative epistatic interactions from systematic patterns of genotype ratio distortions in experimental crosses and wild samples. We then compared the positions of putative incompatibility loci to regions of high genetic differentiation between sympatric sister species as well as between members of clades that may have arisen at the start of this radiation, and further determined if the loci showed fixed differences between the closest living relatives of the lineages ancestral to the hybrid progenitor. Overall, we find little evidence for a major role of intrinsic postzygotic incompatibilities in the Lake Victoria radiation. However, we find putative incompatibility loci significantly more often coinciding with islands of genetic differentiation between species that separated early in the radiation than between the youngest sister species, consistent with the hypothesis that such variants segregated in the hybrid swarm and were sorted between species in the early speciation events.

## Introduction

How and to what extent reproductive isolation emerges during speciation and how it is maintained in the absence of geographic barriers is a fundamental question in evolutionary biology. Postzygotic incompatibilities are factors that reduce the survival or reproduction of hybrid offspring and can hence constitute a barrier to gene flow between populations or species (Coyne & Orr, 2004). Postzygotic incompatibilities can on the one hand arise as a consequence of divergent ecological adaptation, which may result in hybrids having intermediate performance in the ecological niche of either parental species. In the absence of an intermediate or alternative environment, this may reduce hybrid fitness (extrinsic postzygotic incompatibilities) (Schluter, 2000; Nosil, 2012). Alternatively, postzygotic incompatibilities can arise as an ecology-independent by-product of evolutionary divergence between populations, rendering hybrids less fit independent of their environment (intrinsic postzygotic incompatibilities) (Coyne & Orr, 2004). In the well-understood Bateson-Dobzhansky-Muller (BDM) model, postzygotic incompatibilities (‘BDMIs’) arise due to negative epistatic interactions between alleles from different genomic backgrounds that may have fixed between populations due to drift, parallel selection, or divergent selection (Bateson, 1909; Dobzhansky, 1937; Muller, 1942). Often, BDMIs may only become apparent in second or later generations of hybrids, when recessive alleles become fully expressed, referred to as hybrid breakdown (Dobzhansky, 1948; Templeton *et al*., 1986).

BDMIs are expected to contribute to reproductive isolation between populations when they have diverged in geographic separation (in allopatry) and then come into secondary contact, as drift and selection (parallel or divergent) will have led to the accumulation and eventually fixation of genetic differences between the two populations; these differences can then manifest as BDMIs in the hybrid offspring, preventing or reducing further gene flow between the populations (Orr, 1996; Turelli *et al*., 2001; Coyne & Orr, 2004). However, BDMIs are unlikely to emerge if species diverge in geographic proximity (in parapatry or sympatry) and in the presence of gene flow, as alleles contributing to BDMIs will be purged by negative selection in both diverging populations (Gavrilets, 2004; Bank *et al*., 2012).

In the East African cichlid radiations, hundreds of reproductively isolated species have evolved within each of the three largest lakes (Tanganyika, Malawi, and Victoria) in close geographic proximity (reviewed in Seehausen, 2015). In the Lake Victoria haplochromine cichlid adaptive radiation alone, approximately 500 species have evolved within the lake in <15,000 years (Seehausen, 1996; Stager & Johnson, 2008; Meier et al. 2023). The co-existence of up to several dozens of closely related species in any one habitat patch in the lake (Fryer & Iles, 1972; Seehausen, 1996; Turner *et al*., 2001; Genner *et al*., 2004) begs the question how they maintain reproductive isolation from one another. Prezygotic reproductive barriers are known to be important in this system: female mate choice based on male nuptial coloration plays a key role in behavioral reproductive isolation (Seehausen *et al*., 1997; Seehausen & Van Alphen, 1998; Haesler & Seehausen, 2005; Selz *et al*., 2014). In Lake Victoria cichlids, divergence in sexual signaling traits may often have led to divergence in association with divergent ecological adaptation, especially along the steep gradient of light-induced ecological conditions along the water depth axis (Seehausen *et al*., 2008). However, it remains difficult to explain how so many different species can coexist right after speciation within the same water depth and the same microhabitat. Even if behavioral mate choice is an important feature in many of them (Seehausen *et al*., 1998), the question remains how it can evolve so rapidly and persist despite some ongoing gene flow (Meier *et al*., 2018), especially since theoretical studies suggest that in sympatry, disruptive natural selection would have to be unrealistically strong to couple mate choice genes to ecological adaptation genes in the face on gene flow (Felsenstein, 1981; van Doorn *et al*., 2004; Butlin *et al*., 2021). Furthermore, (partial) prezygotic isolation alone may not be effective without any postzygotic isolation (Irwin, 2020; Irwin & Schluter, 2022).

Recent studies have shown that, while all Lake Victoria cichlid species are very young (Meier *et al*., 2017a; 2023; McGee *et al*., 2020), their genomes are mosaics containing much older genetic variants from at least two ancestral lineages (Meier et al. 2023). The modern Lake Victoria radiation evolved within 16,000 years from a hybrid population that arose from three different refugial lineages that had survived the total desiccation of the lake in headwater swamps. All three refugial lineages themselves were leftovers of a hybrid lineage that arose several hundred thousand years ago when two or three deeply divergent lineages met and merged. This hybrid ancestry fueled the radiation by providing large amounts of old genetic variation that could then be sorted into many new combinations. The divergence time of several million years between the ancestral hybridizing lineages that seeded the Lake Victoria region superflock (LVRS) falls into the age range in which we would expect incompatibilities between them to segregate (Stelkens *et al*., 2010; Meier *et al*., 2017a). Indeed, patterns consistent with differential sorting of BDMIs in the LVRS were found: sites that were fixed for alternative alleles in the two ancestral lineages were enriched for species differentiation outlier loci in Lake Victoria (Meier *et al*., 2017a). In the initial hybrid swarm (i.e., a mixture of genomes from at least two divergent ancestral lineages) that seeded the Lake Victoria radiation, BDMIs may thus have been present at high frequencies. On the one hand, the hybrid population would have suffered fitness losses from the presence of BDMIs; on the other hand, they carried large amounts of genetic variation at ecologically relevant loci (and at loci that affect mating traits and mating preferences), which allowed diversification into many different niches. If BDMIs then became coupled to such polymorphisms maintained by negative frequency-dependent or divergent ecological selection, this could have facilitated rapid, repeated and sympatry-robust speciation (Seehausen, 2013). This reorganization of genomic features and incompatibilities from ancestral lineages into different combinations may facilitate rapid evolution of reproductive isolation between the emerging species from a hybrid swarm and their ancestors (Schumer *et al*., 2015; Brennan *et al*., 2019; Marques *et al*., 2019; Wang *et al*., 2021). We thus hypothesized that BDMIs might play a role in speciation and adaptive radiation with gene flow from a hybrid population. If intrinsic postzygotic incompatibilities contribute to reproductive isolation among species of the Lake Victoria radiation, unfavorable combinations of alleles should have been removed by natural selection and should thus be underrepresented among different species. If incompatibilities are sufficiently strong to affect viability of early life stages, they should also be underrepresented among hybrids in experimental species crosses in the lab.

Here, we combine four complementary genomic approaches and datasets to search for signatures consistent with the presence of intrinsic postzygotic incompatibilities (BDMIs) in the Lake Victoria radiation. First, we study segregation patterns in three experimental laboratory F2 crosses between different Lake Victoria cichlid species using restriction site-associated DNA (RAD) marker sequencing to scan hybrid genomes for regions displaying segregation distortion. If there are alleles in the grandparents that do not work well together and cause early life stage mortality when combined in the hybrids, the absence of the genotypes combining these alleles will result in distorted genotype ratios. We also test if there is selection for increased heterozygosity in the F2 hybrids, as predicted under Fisher’s geometric model if the two parental lineages feature co-adapted alleles/genes (Simon *et al*., 2018). Second, we use whole genome sequences from 94 species representing nearly all ecological guilds in the Lake Victoria radiation (1 genome per species) to screen each pair of alleles at loci on different chromosomes for signatures of high linkage disequilibrium among these 94 species (‘multispecies LD’). Unfavorable allelic combinations should have been removed by natural selection and should thus be underrepresented among the different species (or the favorable combinations overrepresented, respectively). Third, we compare the position of putative incompatibility loci relative to highly differentiated regions (based on FST outlier analyses) between sympatric sister species pairs. For the latter we use whole genome sequences of 11 different species pairs with 3-5 individuals per species). We also compare the position of putative incompatibility loci relative to FST outliers between non-sister species representing different clades in the radiation to test if BDMIs may have been sorted in the earlier speciation events when those clades arose (Meier et al. 2023). Finally, we test if putative incompatibility loci correspond to fixed differences in the closest living relatives of the lineages ancestral to the original hybrid swarm at the base of the LVRS.

## Materials and methods

### RAD sequencing data from second-generation (F2) hybrid crosses

The three second-generation (F2) hybrid crosses were between *Pundamilia* sp. ‘nyererei-like’ and *P.* sp. ‘pundamilia-like’, between *P. pundamilia* and *P.* sp. ‘red-head’, and between *P.* sp. ‘nyererei-like’ and *Neochromis omnicaeruleus*. See Feulner *et al*. (2018), Feller *et al*. (2020, 2021), and Feller & Seehausen (2022) for details on breeding, rearing, and processing protocols. The sequencing and genotyping procedures for all three crosses are described in full elsewhere (Feller *et al*., 2021; Feller & Seehausen, 2022). In brief, we generated Restriction-site Associated DNA sequencing (RADseq) libraries that were single-end sequenced (100-150 bp) on an Illumina HiSeq2500 machine. The reads were demultiplexed, quality-filtered, aligned to the anchored version of the *P. nyererei* reference genome (Feulner *et al*., 2018), and variant calling / genotyping was done separately for each cross.

We filtered the three resulting Variant Call Format (VCF) tables using BCFtools (samtools/1.9; Li *et al*., 2009). We only kept bi-allelic SNPs with <50% missing data (genotypes with depth of <10 or quality <20 were set to missing), a minor allele frequency (MAF) of >0.05, and a maximum sequencing depth of less than 1.5 times the interquartile range from the mean. Individuals with >50% missing data, a mean depth of <10, or indications of high amounts of PCR duplication were excluded in the process. We then subset these sets of quality-filtered SNPs to sites that were alternatively homozygous fixed in the grandparents (P) and heterozygous in all first-generation (F1) hybrid parents (‘fixed sites’; note that some P and F1 individuals were not available, see Table S1). Finally, we used Beagle 5.2 (Browning *et al*., 2018; settings: ne=3000, window=10.0) to impute genotypes for the second-generation hybrids (F2s) at all fixed sites, excluding unmapped scaffolds. Table S1 shows an overview of the numbers of SNPs and individuals per cross. Base sequencing depth and quality were high in all three sets (DP >500, QUAL >998).

### Testing for increased heterozygosity in the F2 hybrids

We used the -–het function in vcftools v.0.1.16 (Danecek *et al*., 2011) to output tables of homozygous genotype counts on a per-individual basis. From this, we calculated the proportion of heterozygous genotype counts (at fixed sites) for each individual in R (R Core Team, 2023). Following the approach of (Simon *et al*., 2018), we used Wilcoxon’s signed rank test (wilcox.test function in R) to test if the distribution of heterozygosity in the F2s was symmetrically distributed around the (Mendelian) null expectation of mu=0.5.

### Screening for regions with segregation distortion in the F2 hybrids

To identify and extract regions of segregation distortion in the F2 hybrids we first thinned the sets of fixed sites to remove SNPs that were closely linked. This was done to avoid multiple counts of the same locus in the subsequent tests of overlaps with other potential incompatibility regions. We used JoinMap 4.0 (van Ooijen, 2006) to identify ‘similar loci’ based on a threshold of 0.975 and then removed these in R, additionally removing one of two loci that were still closer than 1,000 bp. This resulted in a final set of 1,066 SNPs for *P*. sp. ‘nyererei-like’ x *P*. sp. ‘pundamilia-like, 921 SNPs for *P. pundamilia* x *P*. sp. ‘red-head’, and 784 SNPs for *P*. sp. ‘nyererei-like’ x *N. omnicaeruleus*.

We then used three complementary approaches to identify segregation distortion. First, we inspected deviations in allele frequencies across all the F2 hybrids in a cross from the Mendelian expectation of 0.5 (‘deviating allele frequency’). We used the –-freq function in vcftools to output per-site allele frequencies. Second, we inspected deviations in genotype ratios from the Mendelian expectation of 1:2:1 (‘segregation distortion’). We used the –-hardy function in vcftools to output genotype counts. In R, we tested each SNP for segregation distortion by applying a Chi-square-test. Then, we subset SNPs with significant segregation distortion (Chi-square test-value of >4.605, i.e., a p-value of <0.1) to those where the genotype ratios approximately conformed to the patterns expected if they were involved in a two-locus (recessive) incompatibility with another SNP. That is, one homozygous genotype should be reduced by 1/16 while the other homozygous genotype and the heterozygous genotypes should each be increased by 1/32. We implemented this in R such that the counts of the less frequent homozygous genotype had to be between 0.6-0.8 times that of the more frequent homozygous genotype, and the counts of the more frequent homozygous genotype had to be between 0.3-0.7 times that of the heterozygous genotypes. Third, we screened for regions with locally increased heterozygosity (‘increased heterozygosity’). Instead of subsetting the SNPs to those conforming to ‘two-locus-incompatibility’-patterns as above, we subsetted them to those with excess heterozygous genotype counts -increased by 10% or more-while both homozygous genotype counts were approximately equally reduced (i.e., the count of one homozygous type had to be min. 0.75 or max. 1.25 times that of the other). For analyses of overlaps with FST outlier windows and high LD pairs (see below), we compiled all three extracted types of distorted regions into one set of distorted regions for each cross.

### Whole genome sequencing data

Sample collection, whole genome re-sequencing, and genotype and variant calling are described in Meier *et al*. (2017a, 2023) and McGee *et al*. (2020). The VCF tables had been filtered by Meier *et al*. (2023) with BCFtools, setting genotypes with a depth of <6 and/or quality of <20 to missing, only keeping bi-allelic SNPS with <50% missing data, and sites were excluded if they overlapped with one of three masks (regions with more than one 35-kmer self-mapping, regions with high repeatability in the reference genome, and regions in which more than 30 of the 400+ genomes had a depth that exceeded the value of mean depth plus 1.5 times the interquartile range); see Meier *et al*. (2023) for more details and used tools. We used a subset of the genomes in these SNP tables and further filtered them as described below.

### Multispecies Inter-Chromosomal Linkage Disequilibrium (LD) analyses across the radiation

To assess non-random allele associations across the species of the Lake Victoria radiation (‘multispecies LD’) we used one male individual of 94 whole genome sequenced Lake Victoria radiation members (Table S2). We first subset the whole genome SNP tables to these 94 individuals plus 13 individuals that represent the closest living relatives to the hybrid swarm ancestors of the Lake Victoria radiation superflock (LVRS) (Meier *et al*., 2017a). The latter include five individuals of *Thoracochromis gracilior* as well as three individuals of *T. pharyngalis* (Nilotic lineage), and four individuals of *Astatotilapia stappersi* as well as one individual of *A.* sp. ‘Yakaema’ (Congolese lineage). We then used BCFtools to filter out sites with >10% missing data and a MAF of <0.05. Next, we subset the SNP tables to the 94 radiation members and only kept sites with <1% missing data, a MAF of >0.05, a depth not exceeding the mean by 1.5 times the interquartile range, and we pruned for intra-chromosomal LD (r^2^>0.9 in 200 bp). This resulted in a set of 464,584 SNPs. We used vcftools to generate the input files for PLINK 1.90 (Chang *et al*., 2015; Purcell & Chang, 2015), which we then used to generate LD statistics for all locus pairs, only outputting pairs with an r^2^-value of >0.2 (settings --r2 ‘inter-chr’ ‘with-freqs’ --ld-window-r2 0.2). We further removed pairs where both SNPs had a MAF of <0.1 and a highly similar ratio between MAF at locus A and MAF at locus B (i.e., a ratio between 0.8-1.2), because locus pairs in high LD with a very low MAF and equal MAFs are most likely to represent a phylogenetic signal. We performed all subsequent analyses on inter-chromosomal pairs with an r^2^-value of >0.5 and with an r^2^-value of >0.6.

### FST outliers

For FST analyses among sympatric sister species as well as among non-sister species of Lake Victoria cichlids we subset the whole genome SNP tables to the 28 species listed in Table S3, for each of which we had genome sequences of two to five male individuals (a total of 108 samples; see Tables S4 and S5 for the list of pairs and numbers of individuals in each). The non-sister species were randomly combined into the same number of pairs as we had for the sympatric sister species, using each species only once. We filtered out sites with >25% missing data, a depth of <10, and a depth exceeding the mean by 1.5 times the interquartile range. This resulted in a set of 47,553,574 SNPs. Within each tested species pair, we then only kept sites with no missing data and a minor allele count of >1, and we calculated Weir and Cockerham’s weighted FST as implemented in vcftools in 50 kb non-overlapping windows. The FST values in windows with less than 10 SNPs were subsequently set to n/a. We chose this approach of using fixed non-overlapping windows because it makes positions of windows directly comparable between all analyzed species pairs and avoids pseudoreplication in outlier detection. (The caveat is that the species pairs will differ in the number and location of missing windows.) For each species pair we identified the top 5% and top 1% outlier windows in R. Pearson’s correlation tests of the weighted FST values were performed to determine if the FST landscapes between species pairs were correlated.

### Overlaying the three datasets

We searched for overlap between SNPs with segregation distortion, SNPs involved in high multi-species LD, and top 5% outlier FST windows, and we tested if the detected overlaps occur more frequently than expected by chance across the genome using a permutation approach. For this, 50kb windows along the genome were assigned two to three different ‘yes/no’ states depending on the tested comparison: 1) is a top 5% FST outlier window, 2) contains a segregation distortion SNP, 3) contains a SNP in high multi-species LD. Before running this analysis, we thinned the SNP sets to a minimum of 50 kb between SNPs, so that if a window contained several SNPs with segregation distortion or high multispecies LD, they were counted as one. (Note that this random thinning is why the r^2^>0.6 set contains some SNPs that are not present in the r^2^>0.5 set in some of the latter analyses.) We first counted the number of observed overlaps, and then randomly shuffled the locations of the ‘yes’ states across the genome 10,000 times, each time counting the generated overlaps. The empirical p-value was calculated as the number of generated counts as or more extreme than the initially observed counts divided by 10’000. We then used Fisher’s method to test if the detected overlaps are more frequent than expected by chance across the 11 species pairs.

### Genotypes of the radiation ancestors at and genes in putative incompatibility regions

We determined if the genotypes of the radiation ancestor’s closest living relatives (Meier *et al*., 2017a) were reciprocally fixed at the SNPs in high multispecies LD, and which of these were also found in top 5% FST outlier windows in any of the 11 sympatric sister species pairs or in a window containing segregation distortion SNPs in either of the three crosses. We then determined which genes contained multiply overlapping and reciprocally fixed SNPs based on the *P. nyererei* v.2 annotation (Feulner *et al*., 2019). Gene names and ontology terms were extracted from Ensembl 109 (Cunningham *et al*., 2022) and UniProt (“UniProt: the Universal Protein knowledgebase in 2023,” 2023). The same analysis was performed for the non-sister species pairs.

The chromosome numbering throughout the manuscript is according to the *P. nyererei* reference genome, to which all alignments were made; see Feulner *et al*. (2018) for the corresponding *Oreochromis niloticus* linkage group numbers.

## Results

### No evidence of selection for increased heterozygosity in F2 hybrids

Mean heterozygosity in the F2 hybrids was not significantly different from the null expectation of 0.5 in two crosses, and even biased towards an overall lower level of heterozygosity in the cross of the very young species pair of P. sp. ‘nyererei-like’ and P. sp. ‘pundamilia-like’ (Meier *et al*., 2017b, 2018) (Table 1).

**Table 1.**
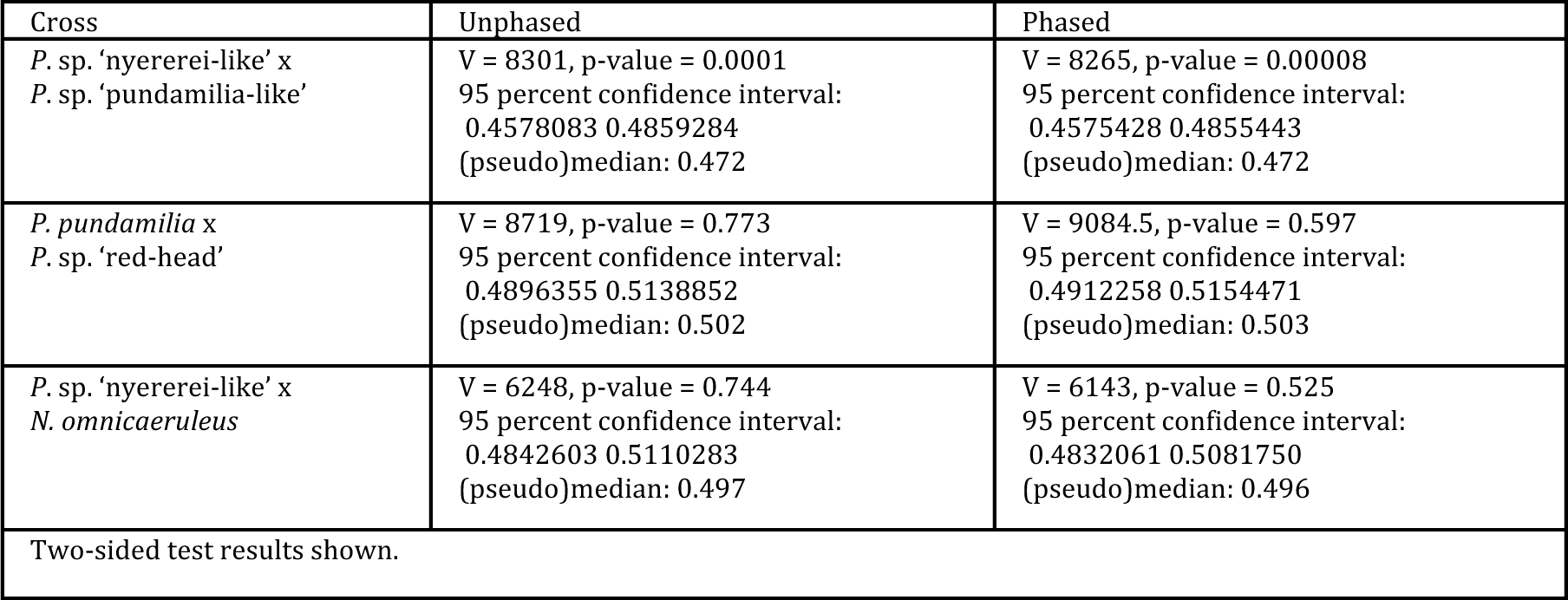
Results of Wilcoxon’s signed rank show that the distribution of heterozygosity in the F2s is not significantly higher than the null expectation of mu=0.5.

### SNPs with segregation distortion in F2 hybrids are distributed across many chromosomes

We detected a total of 116 SNPs with segregation distortion on 15 different chromosomes in *P.* sp. ‘nyererei-like’ x *P.* sp. ‘pundamilia-like’, a total of 64 SNPs on 14 chromosomes in *P. pundam*ilia x P. sp. ‘red-head’, and a total of 27 SNPs on 12 chromosomes in P. sp. ‘nyererei-like’ x N. omnicaeruleus (Fig. 1). A large proportion of the distorted SNPs were found on chromosomes that are implicated in sex determination in some of the species used in the crosses. Chromosome 14 carries a dominant female determiner in *P.* sp. ‘nyererei-like’ (Feller *et al*., 2021), chromosome 10 a dominant male-determiner in *P.* sp. ‘red-head’ (Feulner *et al*., 2018; Feller *et al*., 2021), and chromosome 6 contains a QTL for sex in *P.* sp. ‘nyererei-like’ x *N. omnicaeruleus* (Feller & Seehausen, 2022). Three SNPs on chromosome 14 were shared by *P.* sp. ‘nyererei-like’ x *P.* sp. ‘pundamilia-like’ and *P. pundamilia* x *P.* sp. ‘red-head’. No other segregation distortion SNPs were shared between any cross pair.

**Fig. 1.**
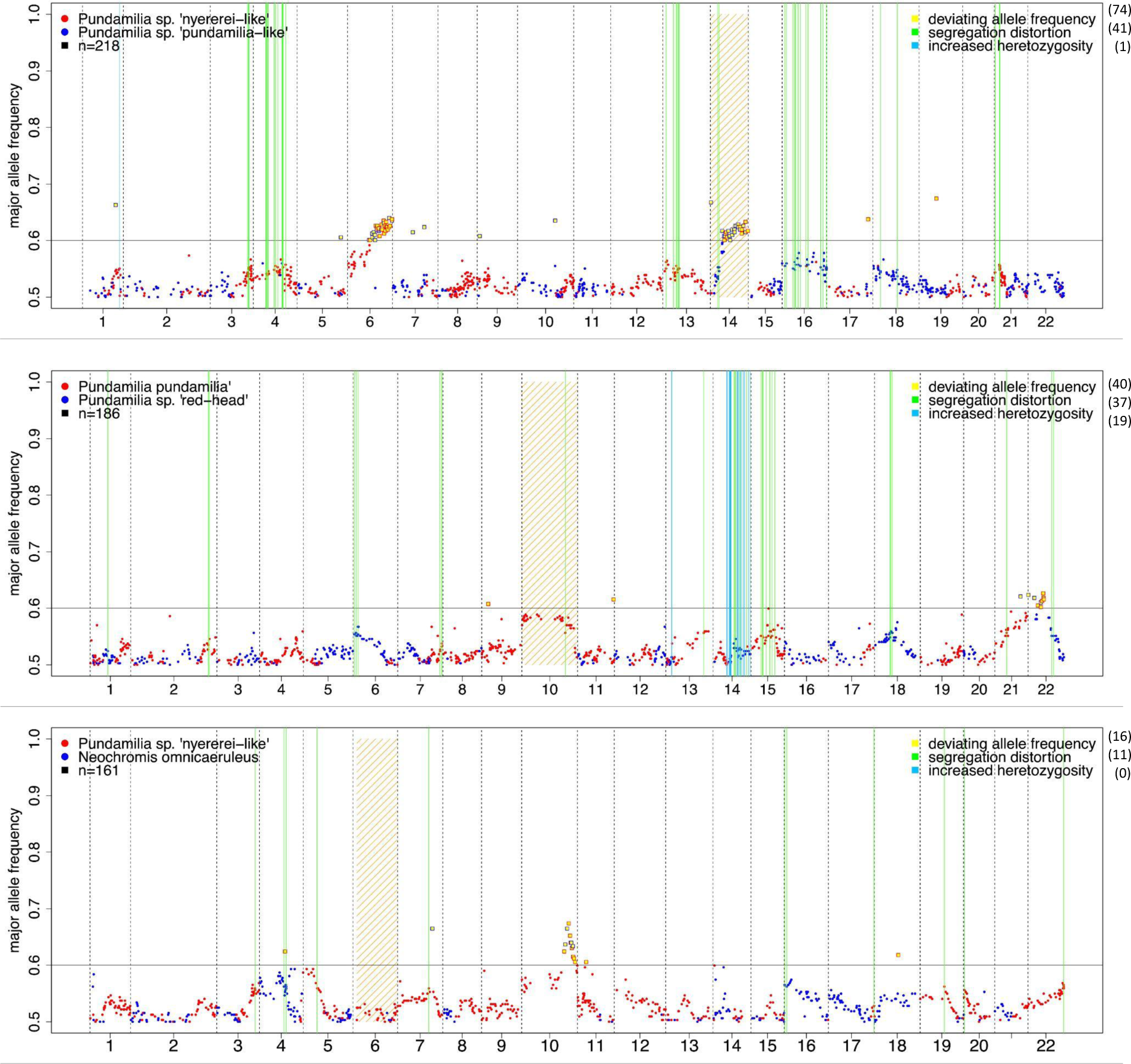
Segregation distortion analyses in three F2 hybrid crosses. The x-axis represents the position in the genome with chromosome numbers according to the *P. nyererei* reference (Feulner *et al*., 2018). The color of the dots indicates which parental species contributed to the more frequent allele at a given SNP. SNPs with a major allele frequency of more than 0.6 (horizontal line) were considered to be deviating in allele frequency (these SNPs are additionally highlighted in yellow with the frame color indicating which parental species contributed the more frequent allele). Green lines indicate the position of SNPs with ‘classic’ segregation distortion (i.e., genotype distortions), and light blue lines indicate the position of SNPs with increased heterozygosity. Chromosomes that are implicated in sex determination in at least one of the cross species are shaded orange. Numbers in brackets are the number of SNPs found in each category (a few appear in more than one category).

### Few overlaps of segregation distortion SNPs with FST outlier windows

Five of the 116 segregation distortion SNPs in *P.* sp. ‘nyererei-like’ x *P.* sp. ‘pundamilia-like’ were located in a 50 kb window containing top 5% FSTs outliers between the two parental species, and one each in *P. pundamilia* x *P.* sp. ‘red-head’ and *P.* sp. ‘nyererei-like’ x *N. omnicaerul*eus (Fig. 2). These overlaps were all not significant (p-values =1).

**Fig. 2.**
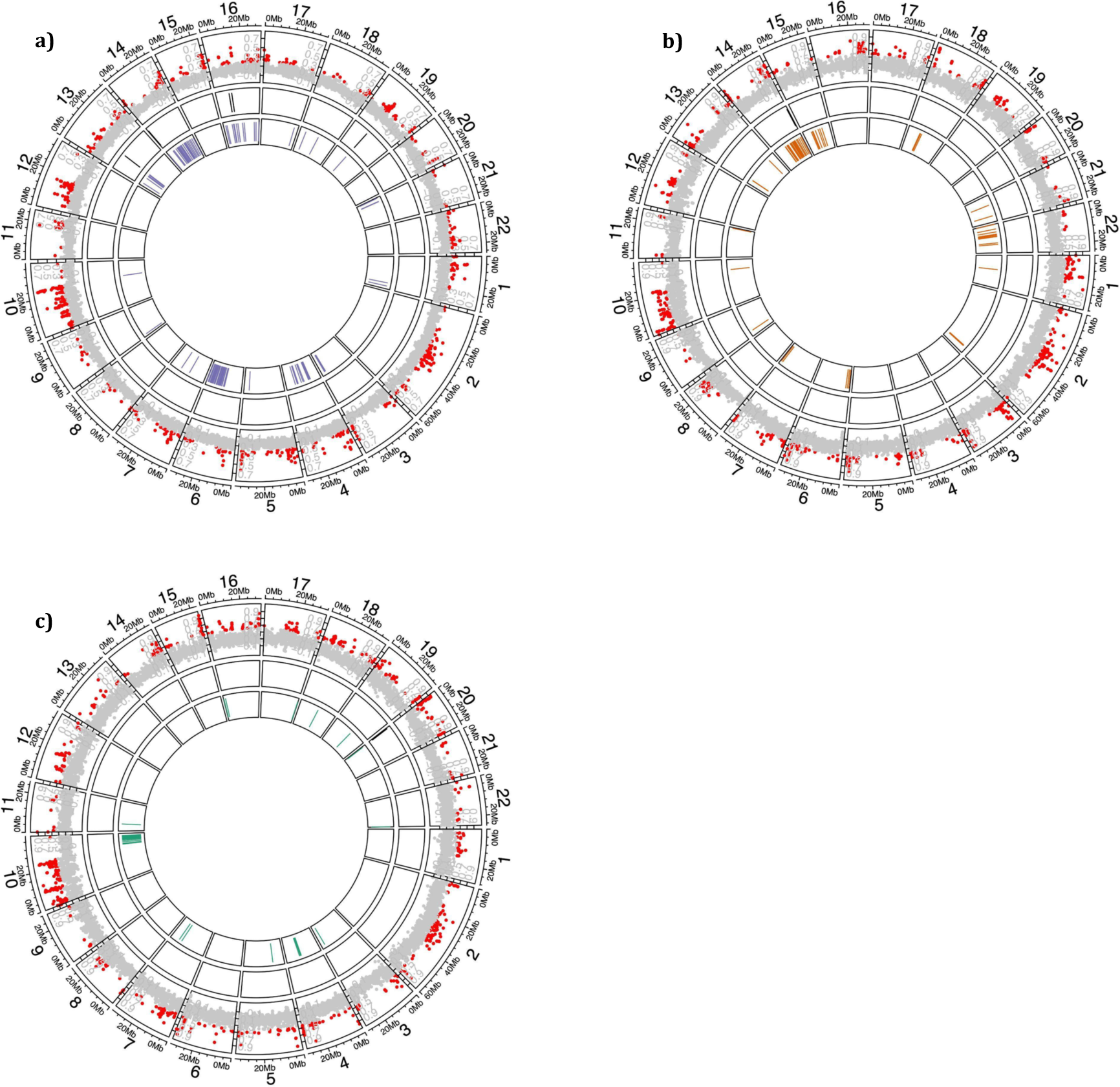
Segregation distortion vs top 5% FST outliers in **a)** is *P*. sp. ‘nyererei-like’ x *P*. sp. ‘pundamilia-like’, **b)** *P. pundamilia* x *P*. sp. ‘red-head’, and **c)** *P*. sp. ‘nyererei-like’ x *N. omnicaeruleus*. Chromosomes are arranged in a circle, with chromosome numbers according to the *P. nyererei* reference. The outer tract shows the FST landscape between the crossed species; top 5% outliers are highlighted red. The inner tract indicates the position of segregation distorted SNPs of all three categories combined (purple for *P*. sp. ‘nyererei-like’ x *P*. sp. ‘pundamilia-like’, orange for *P. pundamilia* x *P*. sp. ‘red-head’, green for *P*. sp. ‘nyererei-like’ x *N. omnicaeruleus*), and the middle tract indicates the positions of 50 kb windows that contained both a top 5% FST outlier and a segregation distortion SNP.

### Strong genome-wide multispecies LD patterns

Of the almost 103 billion inter-chromosomal SNP pairs in the dataset consisting of one genome from each of 94 Lake Victoria haplochromine cichlid species, over 21 million pairs had an r^2^-value of >0.2 and this involved every chromosome (Table 2, Figs. 3, S1). Many thousands of pairs had an even higher r^2^-value, with the count falling below one thousand only at an r^2^-value of >0.6. While these numbers might seem high at a first glance, the percentage of inter-chromosomal pairs with an r^2^>0.2 is only 0.02% (Table 2). Further analyses were performed with the r^2^>0.5 and r^2^>0.6 sets only. Such high r^2^-values can be observed only when multispecies LD between a pair of loci involves many different species across the radiation. The 7,181 SNP pairs with an r^2^>0.5 involved 3,803 ‘unique’ SNPs; unique meaning that SNPs that were linked to multiple other SNPs were only counted once (10,559 SNPs from a total of 2 x 7,181 = 14,362 SNPs were linked to multiple other SNPs). The 625 SNP pairs (i.e., 1250 SNPs) with an r^2^>0.6, involved 602 unique SNPs (648 were linked to multiple other SNPs).

**Fig. 3.**
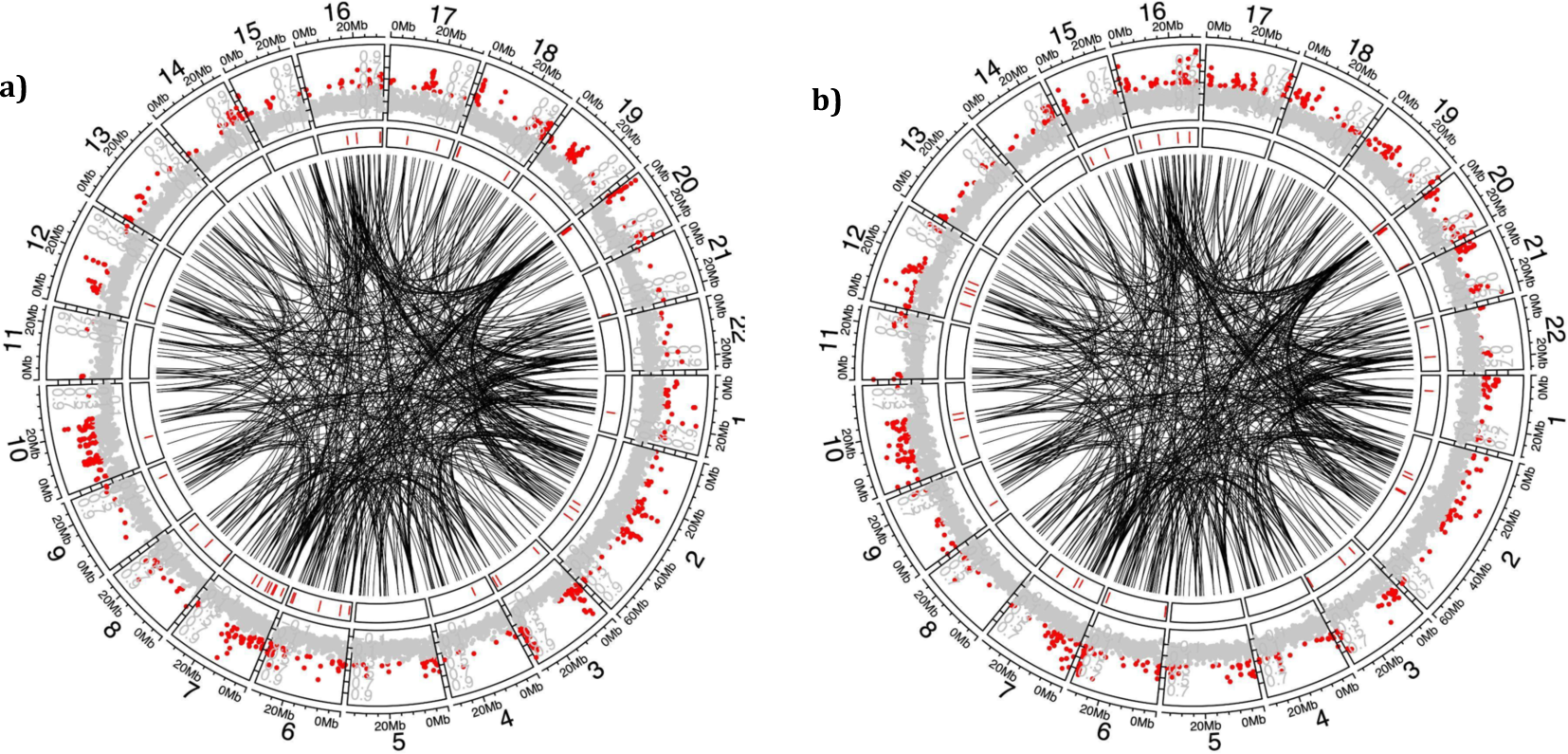
The two sister species pairs in which there were more 50 kb windows than expected by chance that contained both a SNP in high inter-chromosomal LD (r^2^>0.6) and top 5% FST outliers. **a)** *Pundamilia nyererei* vs *Pundamilia pundamilia,* **b)** *Enterochromis antleter* vs *Platytaeniodus* sp. ‘new degeni’. The outer tract shows the FST landscape (gray and red dots; red indicates top 5% outlier windows), and the inner tract (red bars) the location of the overlaps with high inter-chromosomal LD SNPs (r^2^>0.6). The black lines in the middle indicate the connections between the SNPs in multispecies inter-chromosomal LD pairs with an r^2^>0.6.

**Table 2.** Overview of the number of SNP pairs and results of the multispecies LD analyses across 94 Lake Victoria cichlid radiation members. Bold indicates the SNP pairs used in subsequent analyses.

|  |  |
| --- | --- |
| number of SNPs | 464,584 |
| number of SNP pairs total | 107,918,914,236 |
| number of SNP pairs inter-chromosomal | 102,818,692,136 |
| number of SNP pairs with $r^2>0.2$ (total) | 23,036,332 (0.0213% of all pairs) |
| number of SNP pairs with $r^2>0.2$ (inter-chromosomal) | 21,442,178 (0.0209% of all inter-chromosomal pairs) |
| number of SNP pairs with $r^2>0.2$ (intra-chromosomal) | 1,594,154 (0.0313% of all intra-chromosomal pairs) |
| after removing pairs in which SNPs had a similar MAF<0.1 | 12'647'739 |
| number of inter-chromosomal SNP pairs with $r^2>0.3$ | 814,451 |
| number of inter-chromosomal SNP pairs with $r^2>0.4$ | 71,467 |
| **number of inter-chromosomal SNP pairs with $r^2>0.5$** | **7,181** |
| **number of inter-chromosomal SNP pairs with $r^2>0.6$** | **625** |
| number of inter-chromosomal SNP pairs with $r^2>0.7$ | 51 |
| number of inter-chromosomal SNP pairs with $r^2>0.8$ | 12 |
| number of inter-chromosomal SNP pairs with $r^2>0.9$ | 7 |

### Little overlap between the locations of SNPs involved in high multispecies LD and regions with segregation distortion

37 windows that contained at least one SNP with segregation distortion in one of our three crosses also contained at least one SNP in high multispecies LD (r^2^>0.5) across the radiation, although this was not more than expected (p-value = 0.157; Table S6). 18 of these overlaps were found in the cross between *P.* sp. ‘nyererei-like’ and *P.* sp. ‘pundamilia-like’, 13 in the cross between *P. pundamilia* and *P.* sp. ‘red-head’, and six in the cross between *P.* sp. ‘nyererei-like’ and *N. omnicaeruleus.* In the r^2^>0.6 subset, there were six such overlaps, two in each cross. Again, the extent of overlap was not more than expected by chance (p-value = 0.6155; Table S6, Fig. S1).

### SNPs in high multispecies LD and FST outliers significantly coincide in two out of eleven sympatric sister species pairs and in eight out of eleven non-sister species pairs

In all comparisons between 11 pairs of closely related sympatric species, we found that several of the top 5% outlier FST windows contained at least one SNP that was in high multispecies LD (r^2^ >0.6) with another SNP across the radiation. In two comparisons, this overlap was significantly more common than expected by chance (corrected p-value <0.05) (Table 3, Fig. 3). Based on Fisher’s combined probability test, this overlap across the 11 species pairs is significantly more common than expected (62.7 ∼ chi-square (df = 22), combined p-value = 0.000009). A total of 170 unique SNPs involved in high LD (r^2^>0.6) with another SNP overlapped with an FST outlier window in at least one of the 11 species pairs, some occurring in multiple pairs (i.e., there were 317 counts of overlaps involving these 170 SNPs), involving every chromosome except chr14. For the r^2^>0.5 set, a total of 671 unique SNPs involved in high LD with another SNP overlapped with an FST outlier window in the 11 pairs combined, some in multiple pairs (i.e., there were 1184 counts of overlaps involving these 671 SNPs), involving every chromosome. FST landscapes were significantly (positively) correlated between species pairs with one exception (*N. omnicaeruleus* vs *N*. sp. ‘unicuspid scraper’ and *Lithochromis* sp. ‘scraper’ vs *L*. sp. ‘yellow chin’), and they are also significantly (negatively) correlated with the recombination landscape inferred from the *P. pundamilia* x *P*. sp. ‘red-head’ cross by (Feulner *et al*., 2018) (Fig. S2, Table S7).

**Table 3.** Results of permutation tests to assess if SNPs in high multispecies LD significantly coincided with top 5% FST outlier windows in 11 sympatric sister species pairs or 11 non-sister species pairs, respectively. Results shown for r^2^ >0.6 (no significant results for r^2^ >0.5 in sympatric sister species pairs, and in only three non-sister species pairs, indicated by a † behind the species names). P-values were corrected for multiple testing using the false discovery rate method.

| Species pair | Mean weighted FST | Number of overlaps | p-value | significance | corrected p-value | significance (corrected p-value) |
| --- | --- | --- | --- | --- | --- | --- |
| **sympatric sister species pairs** |  |  |  |  |  |  |
| *Pundamilia nyererei* vs *Pundamilia pundamilia* | 0.12 | 47 | 0.000 | *** | 0.000 | *** |
| *Pundamilia* sp. 'nyererei-like' vs *Pundamilia* sp. 'pundamilia-like' | 0.05 | 21 | 0.681 |  | 0.681 |  |
| *Neochromis omnicaruleus* vs *Neochromis* sp. 'unicuspid scraper' | 0.02 | 27 | 0.186 |  | 0.341 |  |
| *Yssichromis pyrrhocephalus* vs *Yssichromis plumbus* | 0.06 | 21 | 0.655 |  | 0.681 |  |
| *Mbipia mbipi* vs *Pundamilia* sp. 'pink anal fin' | 0.04 | 30 | 0.080 | . | 0.220 |  |
| *Neochromis gigas* vs *Paralabidochromis cyaneus* | 0.16 | 27 | 0.182 |  | 0.341 |  |
| *Lithochromis* sp. 'scraper' vs. *Lithochromis* sp. 'yellow chin' | 0.07 | 26 | 0.260 |  | 0.409 |  |
| *Enterochromis I cinctus* (E) vs *Enterochromis II coprologus* (F blue) | 0.09 | 22 | 0.606 |  | 0.681 |  |
| *Enterochromis I* sp. 'new invasive' vs *Yssichromis pyrrhocephalus* | 0.04 | 25 | 0.324 |  | 0.446 |  |
| *Enterochromis I paropius* vs *Enterochromis I coprologus* | 0.04 | 32 | 0.033 | * | 0.121 |  |
| *Enterochromis I antleteri* vs *Platytaeniodus* sp. 'new degeni' | 0.05 | 39 | 0.001 | *** | 0.005 | ** |
| **non-sister species pairs** |  |  |  |  |  |  |
| *Pundamilia nyererei* vs *Neochromis omnicaruleus* † | 0.13 | 48 | 0.000 | *** | 0.000 | *** |
| *Pundamilia pundamilia* vs *Yssichromis pyrrhocephalus* | 0.14 | 32 | 0.031 | * | 0.043 | * |
| *Yssichromis plumbus* vs. *Neochromis gigas* | 0.23 | 25 | 0.334 |  | 0.408 |  |
| *Pundamilia* sp. 'nyererei-like' vs *Lithochromis* sp. 'scraper' † | 0.13 | 52 | 0.000 | *** | 0.000 | *** |
| *Lithochromis* sp. 'yellow chin' vs *Enterochromis I cinctus* (St. E) | 0.07 | 20 | 0.755 |  | 0.755 |  |
| *Pundamilia* sp. 'pundamilia-like' vs *Enterochromis II coprologus* (F blue) | 0.12 | 40 | 0.001 | *** | 0.002 | ** |
| *Mbipia mbipi* vs *Enterochromis I paropius* | 0.08 | 42 | 0.001 | *** | 0.002 | ** |
| *Enterochromis I antleteri* vs *Neochromis* sp. 'unicuspid scraper' | 0.09 | 39 | 0.001 | *** | 0.002 | ** |
| *Paralabidochromis cyaneus* vs *Pundamilia* sp. 'pink anal fin' | 0.14 | 24 | 0.421 |  | 0.463 |  |
| *Enterochromis I* sp. 'new invasive' vs *Pundamilia* sp. 'pundamilia-like' † | 0.15 | 40 | 0.000 | *** | 0.000 | *** |
| *Platytaeniodus* sp. 'new degeni' vs *Pundamilia* sp. 'red head' | 0.13 | 34 | 0.013 | * | 0.02 | * |
| p<0.1, * p<0.05, ** p<0.01, *** p<0.001 |  |  |  |  |  |  |

In eight out of the 11 randomly combined non-sister pairs, the overlap between top 5% FST outlier windows with high multispecies LD SNPs was significant (Table 3). Here too, the overlap across the 11 species pairs was significantly more common than expected (135.249 ∼ chi-square (df = 22), combined p-value= 2.6e-18). We found a similar number of unique and multiple overlapping SNPs as in the sister comparisons (159 unique SNPs at r^2^>0.6 involved in 397 overlaps, and 623 unique SNPs at r^2^>0.5 involved in 1293 overlaps; both distributed across all chromosomes). The majority of the unique high multispecies LD SNPs were the same in the two comparisons (122 out of 170 (sister) and 159 (non-sister) at r^2^>0.6, 456 out of 671 (sister) and 623 (non-sister) at r^2^>0.5). However, the two classes, sister-species and non-sister species differ significantly in the prevalence of having more LD-SNPs coincide with top 5% FST windows (more in the non-sister species, unpaired t-test, one tailed, unequal variances, p=0.0042).

### Few SNPs in high multispecies LD are fixed between the closest living relatives of the hybrid swarm ancestors

Of the 602 unique SNPs in high LD (r^2^>0.6) across the radiation, 22 (∼3.5%) appeared as fixed differences between extant relatives of the radiation ancestors (i.e., five Congolese vs eight Nilotic genomes). The numbers were similar for the r^2^>0.5 SNPs set; 183 SNPs (∼5%) were fixed between the Congolese vs eight Nilotic individuals.

Of the 22 fixed high LD (r^2^>0.6) SNPs, 11 were also located in 50 kb windows with top 5% FST outliers from any of the sympatric sister species pairs, but none were in a window with segregation distortion in our experimental crosses. Of the 183 fixed high LD (r^2^>0.5) SNPs, 32 were located in high FST windows from any of the sympatric sister species pairs, but none were in a window with segregation distortion. No 50kb windows contained all four categories of putatively incompatible SNPs (high LD + fixed differences between ancestors + high FST + segregation distortion). In the non-sister species comparisons, 8 of the 22 fixed high LD (r^2^>0.6) SNPs and 29 of the 183 fixed high LD (r^2^>0.5) SNPs were also located in 50 kb windows with top 5% FST outliers from any of the pairs.

### The majority of putative incompatibility SNPs are found in genic regions

Of the total 40 unique SNPs that are fixed between the ancestral lineages and that are also in high LD among species of the radiation (r^2^>0.5 and/or 0.6) as well as being located in FST outlier windows between sympatric sister species, 25 were located in 25 annotated genes based on the *Pundamilia nyererei* v2 annotation (Feulner *et al*., 2019; Table S8). Of the 36 such SNPs from the non-sister species comparisons (of which 26 were the same SNPs as in the sympatric sister species comparisons), 19 were located in 19 annotated genes (of which 15 were already identified in the sympatric sister species comparisons).

The 29 genes included a variety of functions, among them two genes involved in metabolic processes (Table S8), one involved in regulation of Bone Morphogenetic Protein (BMP) signaling pathway, and the red sensitive opsin gene, *LWS*.

## Discussion

Bateson-Dobzhansky-Muller Incompatibilities (BDMIs) are a frequent cause of reproductive incompatibility between species (Orr, 1996; Turelli *et al*., 2001; Coyne & Orr, 2004). However, the conditions under which BDMIs can establish are limited. BDMIs are unlikely to emerge if species diverge in geographic proximity where gene flow is possible, as alleles involved in BDMIs will be purged by negative selection in both diverging populations (Gavrilets, 2004; Bank *et al*., 2012). It has, however, been proposed that old genetic variants that fixed between species in geographical isolation and act like BDMIs when these species meet and hybridize, could play a role in hybrid speciation (Schumer *et al*., 2015) and adaptive radiation from a hybrid population by becoming rearranged and sorted between populations in the process of speciation (Seehausen, 2013). We experimentally tested this latter hypothesis in the Lake Victoria haplochromine cichlid radiation by screening for underrepresented allele combinations across the genomes of 94 species of the radiation as well as in three experimental crosses between Lake Victoria cichlid species, bred and raised under laboratory conditions.

We did not find evidence for the increased genome-wide heterozygosity in the F2 hybrids that would be expected if the parental species of our three crosses featured many intrinsically co-adapted alleles spread out across the genome (Simon *et al*., 2018; Thompson *et al*., 2022). This suggests that incompatibilities between the parental species are not so ubiquitous that they would induce genome-wide selection for increased heterozygosity (Table 1). This might not be surprising given the young age of this cichlid radiation and that the levels of fixation among species are rather low (Meier *et al*., 2017b, 2018). It is also consistent with the lack of evidence for hybrid breakdown among Lake Victoria cichlid species (Stelkens *et al*., 2015). The observed downward bias in *P*. sp. ‘nyererei-like’ vs *P*. sp. ‘pundamilia-like’ should probably not be over-interpreted. It might have been caused by the fact that only the grandmother and the F1 parents of this cross were available to subset the homozygous fixed SNPs. Hence, not all of the included SNPs may have been truly fixed between the parents, and differing genotype combinations would generate different proportions of heterozygosity levels among the F2s.

Limited by the number of individuals in each F2 cross, we had only modest power to detect the subtle deviations in genotype distortions that would signal the likely presence of incompatibilities. Thus, our goal was to test for patterns consistent with the existence of incompatibilities and not to detect all possible incompatibilities. We detected several regions with segregation distortion in each cross involving many chromosomes (Fig. 1). Almost all detected distorted SNPs were private to one cross, which would not be unexpected in a scenario of sorting of incompatibilities during adaptive radiation from a hybrid population, in which each new emerging species might purge different allele complements involved in incompatibilities (Seehausen, 2013). However, low power to detect regions may also explain low overlap between our crosses. When most incompatibilities become divergently sorted between the first species that arise, fewer should be left to segregate and become divergently fixed in successively later speciation events (Seehausen, 2013) However, to the extent that radiation involves cycles of hybridization, including between non-sister species, and diversification, as in this case (Meier *et al*., 2017b, 2018), some incompatibilities may still become divergently sorted even in recent speciation events. This may be one reason why we do not see more segregation distorted regions in our cross between the more divergent species (*P*. sp. ‘nyererei-like’ x *N. omnicaeruleus*) compared to the two other crosses between more recently diverged species. Another factor to keep in mind is that a large proportion of segregation distortion SNPs are on chromosomes that are known to carry a sex-determining region in at least one of the species used for each cross (Feller *et al*., 2021). Here, segregation distortion could result because sex-linked loci are always heterozygous in the heterogametic sex.

Looking across the radiation with 94 species genomes, we detected a large number of SNPs / SNP pairs in high multispecies LD. This may seem surprising at a first glance, especially when comparing percentages of inter-chromosomal and intrachromosomal pairs with an r^2^-value of >0.2 (Table 2), and in comparison to other studies that have used inter-chromosomal LD to screen for incompatibilities (e.g., Payseur & Place, 2007; Corbett-Detig *et al*., 2013; Schumer *et al*., 2014; but see Hohenlohe *et al*., 2012). However, the number of SNP pairs analyzed here is unprecedented. Furthermore, while the aforementioned studies measured population-level LD among and within a few species, here we measured between-species LD across many species and thus, direct comparisons are not trivial to make. The vast majority of pairs in our case have r^2^<0.2. One explanation for why the percentage of intrachromosomal pairs with r^2^>0.2 is not much higher than that of inter-chromosomal pairs might be that not much physical linkage within chromosomes has been preserved in this radiation due to many generations of recombination since the formation of the ancestral hybrid population (Meier *et al*., 2017a). That is, effective population size in the hybrid population and the emergent radiation was likely high. What needs to be considered though, is that several distinct scenarios could generate the same or similar patterns of high inter-chromosomal LD, and that the presence of intrinsic postzygotic incompatibilities is only one of them. By using one individual per species, we avoided including effects of population structure and species structure (i.e., speciation) on the LD measure, and thus, any LD we detected in our analysis should be due to non-random allele associations shared between multiple species. However, effects of shared evolutionary history between species (i.e., ‘phylogenetic structure’) as well as ecological selection shared between species with similar ecologies are currently not accounted for in our analysis. Effects of phylogenetic structure could reflect shared BDMIs that were sorted in the speciation event that led to the common ancestor of a clade. Any divergent fixation by drift or selection during the origin of the ancestor would result in shared LD between clade members and species in other clades. SNP pairs involving SNPs with a MAF <0.1 and with highly similar MAFs are especially likely to represent effects of such shared phylogenetic history, which is why we additionally excluded such SNP pairs for subsequent analyses. Because there is only very shallow phylogenetic structure among the cichlid species of the Lake Victoria radiation (Bezault *et al*., 2011; McGee *et al*., 2020; Meier et al. 2023), we do not expect large effects on our estimates of inter-chromosomal LD. Effects of shared ecological selection on the other hand, are also possible: Lake Victoria cichlid species fall into many distinct ecological guilds (Table S2; Greenwood, 1974; Witte & Van Oijen, 1990; Seehausen, 1996), and alleles at ecologically and ecomorphologically relevant genes that differentially fixed under divergent ecological selection between members of different guilds might introduce inter-chromosomal LD shared between several species. However, while this is true when the genetic basis for parallel adaptation is simple and non-redundant, it may not be expected with redundancy in the genetic architecture of ecomorphological adaptation. In the latter case, inter-chromosomal LD shared between several species is expected only when the species in a guild share a common ancestor, in which case this is the phylogenetic effect discussed above.

One interesting observation is that in our study, many SNPs are involved in more than one high multispecies LD pair, which could be indicative of higher order epistatic interactions. Incompatibilities might arise from the combination of incompatible (derived) alleles at more than two loci (Satokangas *et al*., 2020) and not all of these may have to be fixed in the parental species (Cutter, 2012). Our screens are thus rather conservative in that they only consider the ‘classic’ two-locus BDMIs. Future investigations should also look at higher order interactions. Other future investigations should consider structural variants such as indels, which was not possible with the current pipeline because it does not call indels reliably (personal comm. D. A. Marques). It is quite likely (McGee *et al*., 2020) that this will add to the number of putative incompatibilities to be discovered.

The analyses we used each have their limitations when considered on their own: modest family sizes for our F2 hybrid crosses meant we had limited power to detect the subtle deviations in genotype ratios we screened for; the multispecies LD analyses may be confounded by phylogenetic structure and effects of ecological adaptation; and FST outliers may not necessarily represent loci or regions relevant for reproductive isolation. However, the strength of our approach lies in the combination of these analyses. Regions that overlap in at least two or in all three analyses are likely to represent incompatibilities that might limit gene flow between species in this largely non-allopatric radiation. This is even more likely if such overlapping SNPs or regions were also divergently fixed between the hybrid swarm ancestors.

There were few overlaps between FST outliers and regions with segregation distortion within crosses (Fig. 2), but several genomic windows contained both a segregation distorted SNP and a SNP in high multispecies LD (r^2^>0.5) (Fig. S1). In two out of the eleven sympatric sister species pairs that we analyzed (three before correction for multiple testing), windows containing a high FST outlier also contained a SNP in high multispecies LD (r^2^>0.5) more often than expected by chance (Table 3, Fig. 3). This was also the case in eight out of eleven tested non-sister species pairs, which would be consistent with the hypothesis that if most incompatibilities become divergently sorted in earlier speciation events, fewer will be left to segregate and become divergently fixed in later speciation events (Seehausen, 2013). About 5% (r^2^>0.5) or 3.5% (r^2^>0.6) of the SNPs in high LD across the radiation were reciprocally fixed between the closest living relatives of the lineages ancestral to the hybrid swarm. 17% (r^2^>0.5) to 50% (r^2^>0.6) of these ancestrally differentially fixed SNPs in high LD were additionally located in a 50kb window containing top 5% FST outliers (from any of the sympatric sister species pairs; similar number for the non-sister pairs). Among the genes at those SNPs are the red-sensitive opsin gene, *LWS*, a gene involved in the regulation of *BMP* signaling, and several genes involved in metabolic and developmental processes (Table S8). These might indeed be good candidates underpinning ecological and reproductive incompatibilities, but further analyses would be needed to determine their importance relative to other genes. LWS is a particularly strong candidate, as it has been implicated in both mate choice and ecological adaptations (Seehausen *et al*., 2008). Although putative incompatibility SNPs that do not map to genes could be playing a regulatory role, genic SNPs seem to be overrepresented among our putative incompatibility SNPs.

It has been proposed that BDMIs may not only contribute to reproductive isolation between populations that come into secondary contact after diverging in allopatry, but they could also play a role in hybrid speciation (Schumer *et al*., 2015) and adaptive radiation and speciation from a hybrid swarm (Seehausen, 2013). Overall, we found modest evidence for a role of intrinsic postzygotic incompatibilities in the Lake Victoria radiation, suggesting that other types of barriers to gene flow were more important in the rapid evolution of the many species in this lake. However, we did find a pattern consistent with the sorting of incompatibilities during speciation from a hybrid swarm where older species pairs have more incompatibilities between them than more recently diverged pairs (Seehausen, 2013). Future work will show if putative incompatibility SNPs are indeed coupled to polymorphisms in ecological or mate choice traits, which could counteract the purging of incompatibilities and facilitate rapid and repeated speciation robust to sympatry.

## Authors’ contributions

**AFF**: Conceptualization, Methodology, Formal analysis, Investigation, Data Curation, Writing - Original Draft, Writing - Review & Editing, Visualization, Project administration;

**CLP**: Conceptualization, Methodology, Writing - Review & Editing, Supervision;

**OS**: Conceptualization (lead), Methodology, Resources, Writing - Review & Editing, Supervision, Project administration, Funding acquisition.

## Acknowledgements

Many thanks to the Seehausen, Peichel, and Hopkins lab groups for many helpful discussions and feedback. Sequence processing and some of the more computationally intensive analyses were performed on a cluster managed by the Genetic Diversity Centre (GDC) at ETH Zurich. This research was funded by Swiss National Science Foundation (SNSF) grant nos. 31003A_163338, 31003A_144046, and 31003A_118293 to OS.

## Data accessibility

Data and code will be available from the Dryad Digital Repository.

