## Supplementary_Tables_and_Figures for "Testing for a role of postzygotic incompatibilities in rapidly speciated Lake Victoria cichlids"

for Feller *et al.*:

**Table S1.** Overview of the number of F2 individuals and SNPs in each cross.

| Cross | **Number of F2 individuals** (males, females) | **Number of quality-filtered SNPs** (of which unmapped scaffolds) | **Number of 'fixed sites'** † excluding unmapped scaffolds |
|---|---|---|---|
| *P.* sp. 'nyererei-like' x *P.* sp. 'pundamilia-like' | 218 (171, 47) | 10,607 (216) | 1,285 |
| *P. pundamilia* x *P.* sp. 'red-head' | 186 (129, 57) | 9,996 (164) | 1,878 |
| *P.* sp. 'nyererei-like' x *N. omnicaeruleus* | 161 (132, 29) | 16,650 (361) | 2,205 |

† Only the female was available for the P of *P.* sp. 'nyererei-like' x *P.* sp. 'pundamilia-like'; only two of the four F1s in *P. pundamilia* x *P.* sp. 'red-head'; only three of the four F1s in *P.* sp. 'nyererei-like' x *N. omnicaeruleus*. See Feller *et al.* (2021) for more information.

**Table S2.** List of samples used in multispecies LD analysis

| Sample | Genus | Species | Location | Sex | Ecology |
|---|---|---|---|---|---|
| 103637 | *Astatotilapia* | *nubila* | Luanso | m | insectivore |
| 11536 | *Astatotilapia* | *nubila* 'rocks' | Python | m | insectivore |
| 109432 | *Astatotilapia* | *nubila* 'swamp red' | Sweya | m | insectivore |
| 104621 | double-stripe group | *tanaos* | MG Transect | m | zooplanktivore |
| 13819 | double-stripe group | *thereuterion* | Makobe | m | insectivore |
| 104013 | *Enterochromis* I | *antleter* blue (St. E) | MG Transect | m | detritivore |
| 104016 | *Enterochromis* I | *cinctus* (St. E) | MG Transect | m | detritivore |
| 108920 | *Enterochromis* I | *coprologus* (St. E) | MG Transect | m | detritivore |
| 103908 | *Enterochromis* I | sp. 'new invasive' | MG Transect | m | detritivore |
| 103720 | *Enterochromis* I | *paropius* | MG Transect | m | detritivore |
| 109795 | *Enterochromis* II | *coprologus* | MG Transect | m | detritivore |
| 11447 | *Enterochromis* II | sp. 'red and blue' | Python | m | detritivore |
| 103767 | *Enterochromis* II | supramacrops 'red tail' | MG Transect | m | detritivore |
| 158683 | *Platytaeniodus* | sp. 'new degeni' | UG3 | m | oral sheller/detritivore |
| TSO4 | *Gaurochromis* | *hiatus* | Sozihe | m | insectivore |
| TMB3 | *Gaurochromis* | *iris* | Magu | m | insectivore |
| 11946 | *Haplochromis* | sp. 'purple yellow' | Makobe | m | epiphytic algae scraper |
| 14561 | *Harpagochromis* | *cavifrons* | Makobe | m | piscivore |
| 161543 | *Harpagochromis* | cf. *pachycephalus* | ? | m | piscivore |
| 109188 | *Harpagochromis* | *howesi* | Makobe | m? | piscivore |
| 78797 | *Harpagochromis* | sp. 'odd dupper' | Anchor | m | piscivore |
| 13135 | *Harpagochromis* | *vonlinnei* | Makobe | m | piscivore |
| 109219 | Incertae sedis | sp. 'large red deepwater' | Makobe | m | insectivore |
| 13269 | Incertae sedis | sp. Lithochrom./Pundam. | Makobe | m | insectivore |
| 14298 | Incertae sedis | sp. 'thick skin' | Makobe | m | insectivore |
| 105005 | Incertae sedis | *vanoijeni* | MG Transect | m | detritivore |
| 80054 | *Labrochromis* | cf. *theliodon* | MG Transect | m | insectivore-pharyngeal-crusher |
| Lishmaeli | *Labrochromis* | *ishmaeli* | captive stock | m | pharyngeal mollusk crusher |
| 14259 | *Labrochromis* | sp. 'stone' | Makobe | m | pharyngeal mollusk crusher |
| 106641 | *Labrochromis* | sp. '1' | MG Transect | m | pharyngeal mollusk crusher |
| 14175 | *Lipochromis* | *cryptodon* | Makobe | m | paedophage |
| 130785 | *Lipochromis* | sp. 'matumbi hunter' | Trawling | m | paedophage |
| 11045 | *Lipochromis* | sp. 'velvet black cryptodon' | Makobe | m | paedophage |
| 80805 | *Lithochromis* | sp. 'brown narrow snout' | Mabibi | m | insectivore |
| 13281 | *Lithochromis* | sp. 'orange' | Makobe | m | insectivore |
| 13393 | *Lithochromis* | sp. 'pseudoblue' | Makobe | m | insectivore |
| 2801 | *Lithochromis* | *rubripinnis* | Luanso | m | insecti-planktivore |
| 11024 | *Lithochromis* | sp. 'scraper' | Makobe | m | epilithic algae scraper |
| 78773 | *Lithochromis* | *xanthopteryx* | Anchor | m | insectivore |
| 109320 | *Lithochromis* | sp. 'yellow chin' | Makobe | m | zooplanktivore |
| 11965 | *Hoplotilapia* | *retrodens* | Makobe | m | oral sheller |
| Ig158 | *Macropleurodus* | *bicolor* | Igombe | m | oral crusher |
| 10448 | *Ptyochromis* | sp. 'deepwater rock sheller' | Python | m | oral sheller |
| 79623 | *Ptyochromis* | *fischeri* | Igombe | m | oral sheller |
| Ig41 | *Ptyochromis* | sp. 'red rock sheller' | Igombe | m? | oral sheller |
| 14245 | *Ptyochromis* | sp. 'striped rock sheller' | Makobe | m | oral sheller |
| 79609 | *Ptyochromis* | *xenognathus* | Igombe | m | oral sheller |
| 10794 | *Ptyochromis* | *xenognathus* 'rocks' | Kissenda | m | oral sheller |
| 11347 | *Mbipia* | *lutea* | Makobe | m | epilithic algae scraper |
| 11003 | *Mbipia* | *mbipi* | Makobe | m | epilithic algae scraper |
| 10791 | *Mbipia* | sp. 'red carp' | Kissenda | m | epilithic algae scraper |
| 11338 | *Neochromis* | *gigas* | Makobe | m | epilithic algae scraper |
| 10897 | *Neochromis* | *greenwoodi* | Anchor | m | epilithic algae scraper |
| Bi10 | *Neochromis* | sp. 'long black' | Bwiru | m | epilithic algae scraper |
| 106816 | *Neochromis* | *omnicaeruleus* | Makobe | m | epilithic algae scraper |
| 10619 | *Neochromis* | *rufocaudalis* | Makobe | m | epilithic algae scraper |
| 11070 | *Neochromis* | sp. 'unicuspid scraper' | Makobe | m | generalist-algae scraper |
| Na15 | *Neochromis* | 'sp. yellow anal scraper' | Nansio | m | epilithic algae scraper |
| 158795 | *Paralabidochromis* | *victoriae* | UG4 | m? | insect picker |
| 10618 | *Paralabidochromis* | *chilotes* | Makobe | m | insectivore |
| 11807 | *Paralabidochromis* | *chromogynos* | Makobe | m | insectivore |
| 130770 | *Paralabidochromis* | *crassilabris* | Sesse | m | insectivore |
| 104042 | *Paralabidochromis* | *plagiodon* | MG Transect | m | oral sheller/detritivore |
| 11639 | *Paralabidochromis* | *plagiodon* 'rocks' | Luanso | m | oral sheller |
| 11049 | *Paralabidochromis* | *sauvagei* | Makobe | m | insectivore |

| | | | | | |
|---|---|---|---|---|---|
| 10628 | *Paralabidochromis* | sp. ‘short snout scraper’ | Makobe | m | epilithic algae scraper |
| 10577 | *Paralabidochromis* | *cyaneus* | Makobe | m | insectivore |
| 80344 | *Paralabidochromis* | *flavus* | Makobe | m | insectivore |
| Ga2 | *Paralabidochromis* | sp. ‘orange anal rockpicker’ | Gana | m | insectivore |
| Ki30 | *Paralabidochromis* | sp. ‘pseudorockpicker’ | Kissenda | m | insectivore |
| Ga33 | *Paralabidochromis* | sp. ‘sky blue picker’ | Gana | m | insectivore |
| 161476 | *Prognathochromis* | dichrourus complex | ? | m | piscivore |
| 130786 | *Prognathochromis* | *perrieri* | lab stock | m | piscivore |
| 10715 | *Pundamilia* | *azurea* | Ruti | m | zooplanktivore |
| 78887 | *Pundamilia* | sp. ‘orange anal nyererei’ | Bihiru | m | insectivore |
| 11034 | *Pundamilia* | sp. ‘pink anal fin’ | Makobe | m | zooplanktivore |
| 84391 | *Pundamilia* | sp. ‘red head’ | ? | m? | insecti-planktivore? |
| E3c | *Pundamilia* | sp. ‘yellow azurea’ | UG/Nsimba | m | insecti-planktivore |
| Ju22 | *Pundamilia* | sp. ‘big blue red’ | Juma | m | insectivore |
| 14125 | *Pundamilia* | sp. ‘deepwater giant’ | Makobe | m | insectivore |
| IG291 | *Pundamilia* | *igneopinnis* | Igombe | m | zooplanktivore |
| 11053 | *Pundamilia* | *nyererei* | Makobe | m | insecti-planktivore |
| 12170 | *Pundamilia* | sp. ‘nyererei-like’ | Python | m | insecti-planktivore |
| 10554 | *Pundamilia* | *pundamilia* | Makobe | m | insectivore |
| 11545 | *Pundamilia* | sp. ‘pundamilia-like’ | Python | m | insectivore |
| 12849 | *Pundamilia* | sp. ‘Luanso red’ | Luanso | m | insectivore |
| 11154 | *Pundamilia* | sp. ‘blue deepwater’ | Luanso | m | insectivore |
| 13653 | *Pundamilia* | *macrocephala* | Makobe | m | insecti-planktivore |
| 12372 | *Pundamilia* | sp. ‘yellow deepwater’ | Luanso | m | zooplanktivore |
| 103778 | *Yssichromis* | cf. *supramacrops* | MG Transect | m | zooplanktivore |
| IG104 | *Yssichromis* | *laparogramma* | Igombe | m | zooplanktivore |
| 79560 | *Yssichromis* | *plumbus* | Igombe | m | zooplanktivore |
| 103754 | *Yssichromis* | *pyrrhocephalus* | MG Transect | m | zooplanktivore |
| Ypiceatus | *Yssichromis* | *piceatus* | lab stock | m? | zooplanktivore |

**Table S3.** List of species used in VCF file used for FST analyses

| **Genus** | **Species** |
|---|---|
| *Enterochromis* I | *paropius* |
| *Enterochromis* I | *antleter* |
| *Enterochromis* I | *cinctus* (St. E) |
| *Enterochromis* I | *coprologus* I |
| *Enterochromis* I | sp. 'new invasive' |
| *Enterochromis* II | *coprologus* F blue morph |
| *Gaurochromis* | *hiatus* |
| *Harpagochromis* | *cavifrons* |
| *Harpagochromis* | *vonlinnei* |
| *Lithochromis* | sp. 'scraper' |
| *Lithochromis* | 'sp. yellow chin' |
| *Mbipia* | *mbipi* |
| *Neochromis* | *gigas* |
| *Neochromis* | *greenwoodi* |
| *Neochromis* | *omnicaeruleus* |
| *Neochromis* | sp. 'unicuspid scraper' |
| *Paralabidochromis* | *sauvagei* |
| *Paralabidochromis* | sp. 'short snout scraper' |
| *Paralabidochromis* | *cyaneus* |
| *Platytaeniodus* | sp. 'new degeni' |
| *Pundamilia* | *nyererei* |
| *Pundamilia* | *pundamilia* |
| *Pundamilia* | sp. 'nyererei-like' |
| *Pundamilia* | sp. 'pundamilia-like' |
| *Pundamilia* | *azurea* |
| *Pundamilia* | sp. 'red head' |
| *Yssichromis* | *pyrrhocephalus* |
| *Yssichromis* | *plumbus* |

| **Table S4.** List of species pairs and number of individuals used in FST analyses - sympatric sister species and cross pairs. | | | | | |
|---|---|---|---|---|---|
| Number of individuals | **Genus** | **Species** | **Location** | **Lake** | **Ecology** |
| **Pair 1** | | | | | |
| 5 | *Pundamilia* | *nyererei* | Makobe | Victoria | insecti-planktivore |
| 5 | *Pundamilia* | *pundamilia* | Makobe | Victoria | insectivore |
| **Pair 2 ('PNP' and 'PPP')** † | | | | | |
| 5 | *Pundamilia* | sp. 'nyererei-like' | Python | Victoria | insecti-planktivore |
| 5 | *Pundamilia* | sp. 'pundamilia-like' | Python | Victoria | insectivore |
| **Pair 3** | | | | | |
| 3 | *Neochromis* | *omnicaeruleus* | Makobe | Victoria | epilithic algae scraper |
| 3 | *Neochromis* | sp. 'unicuspid scraper' | Makobe | Victoria | generalist-algae scraper |
| **Pair 4** | | | | | |
| 3 | *Yssichromis* | *pyrrhocephalus* | MG Transect? | Victoria | zooplanktivore |
| 3 | *Yssichromis* | *plumbus* | ? | Victoria | zooplanktivore |
| **Pair 5** | | | | | |
| 5 | *Mbipia* | *mbipi* | Makobe | Victoria | epilithic algae scraper |
| 5 | *Pundamilia* | sp. 'pink anal fin' | Makobe | Victoria | zooplanktivore |
| **Pair 6** | | | | | |
| 3 | *Neochromis* | *gigas* | Makobe | Victoria | epilithic algae scraper |
| 3 | *Paralabidochromis* | *cyaneus* | Makobe | Victoria | insectivore |
| **Pair 7** | | | | | |
| 3 | *Lithochromis* | sp. 'scraper' | Makobe | Victoria | epilithic algae scraper |
| 3 | *Lithochromis* | sp. 'yellow chin' | Makobe | Victoria | zooplanktivore |
| **Pair 8** | | | | | |
| 5 | *Enterochromis* I | *cinctus* (St. E) | MG Transect | Victoria | detritivore |
| 5 | *Enterochromis* II | *coprologus* F blue morph | MG Transect | Victoria | detritivore |
| **Pair 9** | | | | | |
| 3 | *Enterochromis* I | sp. 'new invasive' | MG Transect | Victoria | detritivore |
| 3 | *Yssichromis* | *pyrrhocephalus* | MG Transect | Victoria | zooplanktivore |
| **Pair 10** | | | | | |
| 4 | *Enterochromis* I | *paropius* | MG Transect | Victoria | detritivore |
| 4 | *Enterochromis* I | *coprologus* I | MG Transect | Victoria | detritivore |
| **Pair 11** | | | | | |
| 3 | *Enterochromis* I | *antleter* | MG Transect | Victoria | detritivore |
| 3 | *Platytaeniodus* | sp. 'new degeni' | MG Transect | Victoria | oral sheller/detritivore |
| **Pair ('PNP' and 'NOM')** † | | | | | |
| 3 | *Pundamilia* | sp. 'nyererei-like' | Python | Victoria | insecti-planktivore |
| 3 | *Neochromis* | *omnicaeruleus* | Makobe | Victoria | epilithic algae scraper |
| **Pair ('PPM' and PRHZ')** † | | | | | |
| 3 | *Pundamilia* | *pundamilia* | Makobe | Victoria | insectivore |
| 3 | *Pundamilia* | sp. 'red head' | Zue? | Victoria | ? |
| † cross species pairs | | | | | |

**Table S5.** List of species pairs and number of individuals used in FST analyses - non-sister species.

| | Genus | Species | Location | Lake | Ecology |
|---|---|---|---|---|---|
| **Pair 1** | | | | | |
| 3 | *Pundamilia* | *nyererei* | Makobe | Victoria | insecti-planktivore |
| 3 | *Neochromis* | *omnicaeruleus* | Makobe | Victoria | epilithic algae scraper |
| **Pair 2** | | | | | |
| 3 | *Pundamilia* | *pundamilia* | Makobe | Victoria | insectivore |
| 3 | *Yssichromis* | *pyrrhocephalus* | MG Transect | Victoria | zooplanktivore |
| **Pair 3** | | | | | |
| 3 | *Yssichromis* | *plumbus* | Igombe | Victoria | zooplanktivore |
| 3 | *Neochromis* | *gigas* | Makobe | Victoria | epilithic algae scraper |
| **Pair 4** | | | | | |
| 3 | *Pundamilia* | sp. 'nyererei-like' | Python | Victoria | insecti-planktivore |
| 3 | *Lithochromis* | sp. 'scraper' | Makobe | Victoria | epilithic algae scraper |
| **Pair 5** | | | | | |
| 3 | *Lithochromis* | sp. 'yellow chin' | Makobe | Victoria | zooplanktivore |
| 3 | *Enterochromis* I | *cinctus* (St. E) | MG Transect | Victoria | detritivore |
| **Pair 6** | | | | | |
| 5 | *Pundamilia* | sp. 'pundamilia-like' | Python | Victoria | insectivore |
| 5 | *Enterochromis* II | *coprologus* F blue morph | MG Transect | Victoria | detritivore |
| **Pair 7** | | | | | |
| 5 | *Mbipia* | *mbipi* | Makobe | Victoria | epilithic algae scraper |
| 5 | *Enterochromis* I | *paropius* | MG Transect | Victoria | detritivore |
| **Pair 8** | | | | | |
| 3 | *Enterochromis* I | *antleter* | MG Transect | Victoria | detritivore |
| 3 | *Neochromis* | sp. 'unicuspid scraper' | Makobe | Victoria | generalist-algae scraper |
| **Pair 9** | | | | | |
| 3 | *Paralabidochromis* | *cyaneus* | Makobe | Victoria | insectivore |
| 3 | *Pundamilia* | sp. 'pink anal fin' | Makobe | Victoria | zooplanktivore |
| **Pair 10** | | | | | |
| 3 | *Enterochromis* I | sp. 'new invasive' | MG Transect | Victoria | detritivore |
| 3 | *Pundamilia* | sp. 'pundamilia-like' | Python | Victoria | insectivore |
| **Pair 11** | | | | | |
| 3 | *Platytaeniodus* | sp. 'new degeni' | MG Transect | Victoria | oral sheller/detritivore |
| 3 | *Pundamilia* | sp. 'red head' | Zue | Victoria | ? |

**Table S6.** Windows containing both a SNP with segregation distortion in either of the three crosses and a SNP in high multispecies LD with an $r^2$ of >0.5. Bold windows are also overlapping in the $r^2$ >0.6 subset and the italic window only appears in the latter subset, which is due to the thinning method where only one of two SNPs within 50 kb of each other were kept (randomly).

| chromosome | start | end | found in cross |
|---|---|---|---|
| chr1 | 12350000 | 12399999 | PPMxPRHZ |
| **chr4** | **20650000** | **20699999** | **PNPxPPP** |
| chr4 | 22400000 | 22449999 | PNPxPPP |
| **chr6** | **850000** | **899999** | **PPMxPRHZ** |
| chr6 | 20700000 | 20749999 | PNPxPPP |
| **chr6** | **21550000** | **21599999** | **PNPxPPP** |
| chr6 | 26000000 | 26049999 | PNPxPPP |
| chr6 | 27300000 | 27349999 | PNPxPPP |
| chr6 | 27900000 | 27949999 | PNPxPPP |
| chr6 | 30900000 | 30949999 | PNPxPPP |
| chr7 | 29500000 | 29549999 | PPMxPRHZ |
| chr9 | 4500000 | 4549999 | PPMxPRHZ |
| chr10 | 29700000 | 29749999 | PNPxNOM |
| chr10 | 34050000 | 34099999 | PNPxNOM |
| chr10 | 37200000 | 37249999 | PNPxNOM |
| chr13 | 11500000 | 11549999 | PNPxPPP |
| chr14 | 350000 | 399999 | PNPxPPP |
| chr14 | 6050000 | 6099999 | PNPxPPP |
| chr14 | 12150000 | 12199999 | PPMxPRHZ |
| chr14 | 12300000 | 12349999 | PPMxPRHZ |
| chr14 | 12450000 | 12499999 | PNPxPPP |
| chr14 | 15600000 | 15649999 | PPMxPRHZ |
| chr14 | 19350000 | 19399999 | PNPxPPP |
| chr14 | 20550000 | 20599999 | PNPxPPP |
| chr14 | 20800000 | 20849999 | PPMxPRHZ |
| chr14 | 22900000 | 22949999 | PNPxPPP |
| chr15 | 6900000 | 6949999 | PPMxPRHZ |
| chr15 | 10550000 | 10599999 | PPMxPRHZ |
| chr16 | 1700000 | 1749999 | PNPxNOM |
| chr16 | 1900000 | 1949999 | PNPxPPP |
| chr16 | 7550000 | 7599999 | PNPxPPP |
| chr16 | 8800000 | 8849999 | PNPxPPP |
| chr18 | 10750000 | 10799999 | PPMxPRHZ |
| *chr18* | *16650000* | *16700000* | *PNPxNOM* |
| **chr20** | **650000** | **699999** | **PNPxNOM** |
| chr22 | 4300000 | 4349999 | PPMxPRHZ |
| **chr22** | **17700000** | **17749999** | **PPMxPRHZ** |
| chr22 | 24950000 | 24999999 | PNPxNOM |

| **Table S7.** Results of Pearson's correlation tests of weighted FST scores between sympatric sister species pairs. | | | | | | | |
|---|---|---|---|---|---|---|---|
| **pair1** | **pair2** | **t** | **df** | **CI low.** | **CI up.** | **corr.** | **p-value** |
| Pundamilia nyererei vs Pundamilia pundamilia | Pundamilia sp. 'nyererei-like vs Pundamilia sp. 'pundamilia-like' | 16.86 | 12672 | 0.131 | 0.165 | 0.148 | 4.2E-63 |
| Pundamilia nyererei vs Pundamilia pundamilia | Neochromis omnicaeruleus vs Neochromis sp. 'unicuspid scraper' | 11.98 | 12536 | 0.089 | 0.124 | 0.106 | 6.5E-33 |
| Pundamilia nyererei vs Pundamilia pundamilia | Yssichromis pyrrhocephalus vs Yssichromis plumbus | 16.71 | 12525 | 0.131 | 0.165 | 0.148 | 4.8E-62 |
| Pundamilia nyererei vs Pundamilia pundamilia | Neochromis gigas vs Paralabidochromis cyaneus | 28.13 | 12735 | 0.225 | 0.258 | 0.242 | 5.6E-169 |
| Pundamilia nyererei vs Pundamilia pundamilia | Neochromis gigas vs Paralabidochromis cyaneus | 12.34 | 12549 | 0.092 | 0.127 | 0.109 | 8.8E-35 |
| Pundamilia nyererei vs Pundamilia pundamilia | Lithochromis sp. 'scraper' vs Lithochromis sp. 'yellow chin' | 14.72 | 12650 | 0.113 | 0.147 | 0.130 | 1.1E-48 |
| Pundamilia nyererei vs Pundamilia pundamilia | Enterochromis I cinctus (E) vs Enterochromis II coprologus (F blue) | 25.83 | 12721 | 0.207 | 0.240 | 0.223 | 2.2E-143 |
| Pundamilia nyererei vs Pundamilia pundamilia | Enterochromis I sp. 'new invasive' vs Yssichromis pyrrhocephalus | 23.38 | 12620 | 0.187 | 0.220 | 0.204 | 2.2E-118 |
| Pundamilia nyererei vs Pundamilia pundamilia | Enterochromis I paropius vs Enterochromis I coprologus | 26.02 | 12711 | 0.208 | 0.241 | 0.225 | 1.7E-145 |
| Pundamilia nyererei vs Pundamilia pundamilia | Enterochromis I antleter vs Platytaeniodus 'new degeni' | 22.91 | 12622 | 0.183 | 0.216 | 0.200 | 8.3E-114 |
| Pundamilia sp. 'nyererei-like vs Pundamilia sp. 'pundamilia-like' | Neochromis omnicaeruleus vs Neochromis sp. 'unicuspid scraper' | 6.52 | 12525 | 0.041 | 0.076 | 0.058 | 7.4E-11 |
| Pundamilia sp. 'nyererei-like vs Pundamilia sp. 'pundamilia-like' | Yssichromis pyrrhocephalus vs Yssichromis plumbus | 8.51 | 12501 | 0.058 | 0.093 | 0.076 | 2.0E-17 |
| Pundamilia sp. 'nyererei-like vs Pundamilia sp. 'pundamilia-like' | Mbipia mbipi vs Pundamilia sp. 'pink anal fin' | 10.18 | 12718 | 0.073 | 0.107 | 0.090 | 2.9E-24 |
| Pundamilia sp. 'nyererei-like vs Pundamilia sp. 'pundamilia-like' | Neochromis gigas vs Paralabidochromis cyaneus | 8.13 | 12537 | 0.055 | 0.090 | 0.072 | 4.7E-16 |
| Pundamilia sp. 'nyererei-like vs Pundamilia sp. 'pundamilia-like' | Lithochromis sp. 'scraper' vs Lithochromis sp. 'yellow chin' | 6.92 | 12625 | 0.044 | 0.079 | 0.061 | 4.7E-12 |
| Pundamilia sp. 'nyererei-like vs Pundamilia sp. 'pundamilia-like' | Enterochromis I cinctus (E) vs Enterochromis II coprologus (F blue) | 15.84 | 12701 | 0.122 | 0.156 | 0.139 | 5.5E-56 |
| Pundamilia sp. 'nyererei-like vs Pundamilia sp. 'pundamilia-like' | Enterochromis I sp. 'new invasive' vs Yssichromis pyrrhocephalus | 12.10 | 12589 | 0.090 | 0.124 | 0.107 | 1.6E-33 |
| Pundamilia sp. 'nyererei-like vs Pundamilia sp. 'pundamilia-like' | Enterochromis I paropius vs Enterochromis I coprologus | 15.45 | 12696 | 0.119 | 0.153 | 0.136 | 2.2E-53 |
| Pundamilia sp. 'nyererei-like vs Pundamilia sp. 'pundamilia-like' | Enterochromis I antleter vs Platytaeniodus sp. 'new degeni' | 11.77 | 12604 | 0.087 | 0.121 | 0.104 | 8.3E-32 |
| Neochromis omnicaeruleus vs Neochromis sp. 'unicuspid scraper' | Yssichromis pyrrhocephalus vs Yssichromis plumbus | 10.61 | 12412 | 0.077 | 0.112 | 0.095 | 3.5E-26 |
| Neochromis omnicaeruleus vs Neochromis sp. 'unicuspid scraper' | Mbipia mbipi vs Pundamilia sp. 'pink anal fin' | 7.19 | 12571 | 0.047 | 0.081 | 0.064 | 7.1E-13 |
| Neochromis omnicaeruleus vs Neochromis sp. 'unicuspid scraper' | Neochromis gigas vs Paralabidochromis cyaneus | 2.18 | 12453 | 0.002 | 0.037 | 0.020 | 2.9E-02 |
| Neochromis omnicaeruleus vs Neochromis sp. 'unicuspid scraper' | Lithochromis sp. 'scraper' vs Lithochromis sp. 'yellow chin' | 1.31 | 12498 | -0.006 | 0.029 | 0.012 | 1.9E-01 |
| Neochromis omnicaeruleus vs Neochromis sp. 'unicuspid scraper' | Enterochromis I cinctus (E) vs Enterochromis II coprologus (F blue) | 10.02 | 12558 | 0.072 | 0.106 | 0.089 | 1.6E-23 |
| Neochromis omnicaeruleus vs Neochromis sp. 'unicuspid scraper' | Enterochromis I sp. 'new invasive' vs Yssichromis pyrrhocephalus | 6.17 | 12481 | 0.038 | 0.073 | 0.055 | 6.9E-10 |
| Neochromis omnicaeruleus vs Neochromis sp. 'unicuspid scraper' | Enterochromis I paropius vs Enterochromis I coprologus | 5.87 | 12552 | 0.035 | 0.070 | 0.052 | 4.4E-09 |
| Neochromis omnicaeruleus vs Neochromis sp. 'unicuspid scraper' | Enterochromis I antleter vs Platytaeniodus sp. 'new degeni' | 5.16 | 12504 | 0.029 | 0.064 | 0.046 | 2.5E-07 |
| Yssichromis pyrrhocephalus vs Yssichromis plumbus | Mbipia mbipi vs Pundamilia sp. 'pink anal fin' | 13.85 | 12574 | 0.105 | 0.140 | 0.123 | 2.8E-43 |
| Yssichromis pyrrhocephalus vs Yssichromis plumbus | Neochromis gigas vs Paralabidochromis cyaneus | 3.94 | 12429 | 0.018 | 0.053 | 0.035 | 8.2E-05 |
| Yssichromis pyrrhocephalus vs Yssichromis plumbus | Lithochromis sp. 'scraper' vs Lithochromis sp. 'yellow chin' | 4.62 | 12488 | 0.024 | 0.059 | 0.041 | 3.8E-06 |
| Yssichromis pyrrhocephalus vs Yssichromis plumbus | Enterochromis I cinctus (E) vs Enterochromis II coprologus (F blue) | 8.78 | 12563 | 0.061 | 0.095 | 0.078 | 1.9E-18 |
| Yssichromis pyrrhocephalus vs Yssichromis plumbus | Enterochromis I sp. 'new invasive' vs Yssichromis pyrrhocephalus | 30.51 | 12547 | 0.246 | 0.279 | 0.263 | 3.0E-197 |
| Yssichromis pyrrhocephalus vs Yssichromis plumbus | Enterochromis I paropius vs Enterochromis I coprologus | 13.13 | 12566 | 0.099 | 0.134 | 0.116 | 3.9E-39 |
| Yssichromis pyrrhocephalus vs Yssichromis plumbus | Enterochromis I antleter vs Platytaeniodus sp. 'new degeni' | 12.07 | 12505 | 0.090 | 0.125 | 0.107 | 2.3E-33 |
| Mbipia mbipi vs Pundamilia sp. 'pink anal fin' | Neochromis gigas vs Paralabidochromis cyaneus | 5.78 | 12588 | 0.034 | 0.069 | 0.051 | 7.8E-09 |
| Mbipia mbipi vs Pundamilia sp. 'pink anal fin' | Lithochromis sp. 'scraper' vs Lithochromis sp. 'yellow chin' | 4.87 | 12701 | 0.026 | 0.060 | 0.043 | 1.2E-06 |
| Mbipia mbipi vs Pundamilia sp. 'pink anal fin' | Enterochromis I cinctus (E) vs Enterochromis II coprologus (F blue) | 11.44 | 12794 | 0.083 | 0.118 | 0.101 | 3.8E-30 |
| Mbipia mbipi vs Pundamilia sp. 'pink anal fin' | Enterochromis I sp. 'new invasive' vs Yssichromis pyrrhocephalus | 14.20 | 12669 | 0.108 | 0.142 | 0.125 | 2.0E-45 |
| Mbipia mbipi vs Pundamilia sp. 'pink anal fin' | Enterochromis I paropius vs Enterochromis I coprologus | 21.34 | 12777 | 0.169 | 0.202 | 0.185 | 2.8E-99 |

| | | | | | | | |
|---|---|---|---|---|---|---|---|
| Mbipia mbipi vs Pundamilia sp. 'pink anal fin' | Enterochromis I antleter vs Platytaeniodus sp. 'new degeni' | 20.22 | 12670 | 0.160 | 0.194 | 0.177 | 1.7E-89 |
| Neochromis gigas vs Paralabidochromis cyaneus | Lithochromis sp. 'scraper' vs Lithochromissp. ' yellow chin' | 6.45 | 12524 | 0.040 | 0.075 | 0.058 | 1.1E-10 |
| Neochromis gigas vs Paralabidochromis cyaneus | Enterochromis I cinctus (E) vs Enterochromis II coprologus (F blue) | 7.06 | 12578 | 0.045 | 0.080 | 0.063 | 1.7E-12 |
| Neochromis gigas vs Paralabidochromis cyaneus | Enterochromis I sp. 'new invasive' vs Yssichromis pyrrhocephalus | 8.47 | 12493 | 0.058 | 0.093 | 0.076 | 2.6E-17 |
| Neochromis gigas vs Paralabidochromis cyaneus | Enterochromis I paropius vs Enterochromis I coprologus | 7.57 | 12583 | 0.050 | 0.085 | 0.067 | 4.0E-14 |
| Neochromis gigas vs Paralabidochromis cyaneus | Enterochromis I antleter vs Platytaeniodus sp. 'new degeni' | 10.62 | 12520 | 0.077 | 0.112 | 0.095 | 3.0E-26 |
| Lithochromis sp. 'scraper' vs Lithochromis sp. 'yellow chin' | Enterochromis I cinctus (E) vs Enterochromis II coprologus (F blue) | 12.06 | 12701 | 0.089 | 0.124 | 0.106 | 2.7E-33 |
| Lithochromis sp. 'scraper' vs Lithochromis sp. 'yellow chin' | Enterochromis I sp. 'new invasive' vs Yssichromis pyrrhocephalus | 11.80 | 12578 | 0.087 | 0.122 | 0.105 | 5.6E-32 |
| Lithochromis sp. 'scraper' vs Lithochromis sp. 'yellow chin' | Enterochromis I paropius vs Enterochromis I coprologus | 12.76 | 12676 | 0.095 | 0.130 | 0.113 | 4.5E-37 |
| Lithochromis sp. 'scraper' vs Lithochromis sp. 'yellow chin' | Enterochromis I antleter vs Platytaeniodus sp. 'new degeni' | 12.02 | 12601 | 0.089 | 0.124 | 0.106 | 4.4E-33 |
| Enterochromis I cinctus (E) vs Enterochromis II coprologus (F blue) | Enterochromis I sp. 'new invasive' vs Yssichromis pyrrhocephalus | 12.40 | 12658 | 0.092 | 0.127 | 0.110 | 4.3E-35 |
| Enterochromis I cinctus (E) vs Enterochromis II coprologus (F blue) | Enterochromis I paropius vs Enterochromis I coprologus | 11.51 | 12772 | 0.084 | 0.118 | 0.101 | 1.7E-30 |
| Enterochromis I cinctus (E) vs Enterochromis II coprologus (F blue) | Enterochromis I antleter vs Platytaeniodus sp. 'new degeni' | 11.78 | 12668 | 0.087 | 0.121 | 0.104 | 6.9E-32 |
| Enterochromis I sp. 'new invasive' vs Yssichromis pyrrhocephalus | Enterochromis I paropius vs Enterochromis I coprologus | 16.85 | 12668 | 0.131 | 0.165 | 0.148 | 4.8E-63 |
| Enterochromis I sp. 'new invasive' vs Yssichromis pyrrhocephalus | Enterochromis I antleter vs Platytaeniodus sp. 'new degeni' | 13.47 | 12583 | 0.102 | 0.136 | 0.119 | 4.7E-41 |
| Enterochromis I paropius vs Enterochromis I coprologus | Enterochromis I antleter vs Platytaeniodus sp. 'new degeni' | 14.94 | 12659 | 0.114 | 0.149 | 0.132 | 5.1E-50 |
| **Results of Pearson's correlation tests of weighted FST scores with recombination rate** | | | | | | | |
| Pundamilia nyererei vs Pundamilia pundamilia | | -6.37 | 1542 | -0.208 | -0.111 | -0.160 | 2.5E-10 |
| Pundamilia sp. 'nyererei-like vs Pundamilia sp. 'pundamilia-like' | | -5.97 | 1538 | -0.199 | -0.101 | -0.151 | 2.9E-09 |
| Neochromis omnicaeruleus vs Neochromis sp. 'unicuspid scraper' | | -2.94 | 1526 | -0.125 | -0.025 | -0.075 | 3.4E-03 |
| Yssichromis pyrrhocephalus vs Yssichromis plumbus | | -4.50 | 1525 | -0.164 | -0.065 | -0.114 | 7.5E-06 |
| Mbipia mbipi vs Pundamilia sp. 'pink anal fin' | | -6.06 | 1545 | -0.201 | -0.103 | -0.152 | 1.7E-09 |
| Neochromis gigas vs Paralabidochromis cyaneus | | -2.86 | 1523 | -0.123 | -0.023 | -0.073 | 4.2E-03 |
| Lithochromis sp. 'scraper' vs Lithochromis sp. 'yellow chin' | | -2.66 | 1549 | -0.117 | -0.018 | -0.067 | 8.0E-03 |
| Enterochromis I cinctus (E) vs Enterochromis II coprologus (F blue) | | -5.69 | 1548 | -0.191 | -0.094 | -0.143 | 1.6E-08 |
| Enterochromis I sp. 'new invasive' vs Yssichromis pyrrhocephalus | | -3.26 | 1536 | -0.132 | -0.033 | -0.083 | 1.1E-03 |
| Enterochromis I paropius vs Enterochromis I coprologus | | -4.44 | 1547 | -0.161 | -0.063 | -0.112 | 9.4E-06 |
| Enterochromis I antleter vs Platytaeniodus sp. 'new degeni' | | -4.29 | 1537 | -0.158 | -0.059 | -0.109 | 1.9E-05 |

**Table S8**. List of genes in which high multispecies LD ($r^2$ >0.5 and/or >0.6) SNPs are located that are fixed between the extant relatives of the radiation ancestors and are also situated in windows with top 5% FST outliers (between any of the tested species pairs).

| chr | start | end | gene name | Makobe island cichlid gene (P. nyererei v.1) on Ensembl | GO annotations (UniProt) cellular component | GO annotations (UniProt) molecular function | GO annotations (UniProt) biological process | sister pairs | non-sister pairs |
|---|---|---|---|---|---|---|---|---|---|
| 1 | 11402815 | 11405602 | mrpl48 | Mitochondrial ribosomal protein L48 | mitochondrial ribosome | n/a | n/a | 3 | 1 |
| 2 | 24101268 | 24236959 | mgat4c | Alpha-1,3 -mannosyl-glycoprotein 4-beta-N-acetyl glucosaminyl transferase C-like | n/a | n/a | n/a | 1,2,4,7,8,9 | 3,4,5,10,11 |
| 2 | 26199951 | 26207444 | dusp11 | Dual specificity phosphatase 11 | n/a | protein tyrosine/ serine/ threonine phosphatase activity | protein dephosphorylation | 4,5 | 7 |
| 2 | 28005791 | 28180551 | cadps2 | Ca++-dependent secretion activator 2 | cytoplasmic vesicle, presynapse | n/a | dense core granule exocytosis, synaptic vesicle exocytosis | 4,5 | 4,5,7,9 |
| 4 | 3663902 | 3801979 | man1a2 | mannosidase alpha class 1A member 2 | membrane | calcium ion binding, mannosyl-oligosaccharide 1,2-alpha-mannosidase activity | carbohydrate metabolic process, determination of heart/ liver/ pancreatic left/right asymmetry, intrahepatic bile duct development, regulation of cilium assembly | n/a | 10 |
| 5 | 316469 | 328106 | si:ch211-112g6.4 | uncharacterized LOC102211607 | n/a | n/a | n/a | 9 | 3 |
| 6 | 1526352 | 1853564 | GRIK2 | glutamate receptor ionotropic, kainate 2-like | postsynaptic membrane | ionotropic glutamate receptor activity | n/a | 3,5,10 | 1,3,6,7 |
| 6 | 5415106 | 5603199 | ptpru | Protein tyrosine phosphatase receptor type Ub | membrane | protein tyrosine phosphatase activity | protein dephosphorylation | 2,8,11 | n/a |
| 6 | 9549914 | 9574871 | zhx2 | zinc fingers and homeoboxes 2 | nucleus | DNA binding | n/a | n/a | 6 |
| 7 | 798190 | 887629 | adgrl2 | Adhesion G protein-coupled receptor L2a | membrane | carbohydrate binding, G protein-coupled receptor activity | cell surface receptor signaling pathway | 8,11 | 2,9 |
| 7 | 4721822 | 4747907 | armc9 | Armadillo repeat containing 9 | centriole, ciliary basal body, cytoplasm | n/a | cilium assembly | 1,8 | n/a |
| 9 | 24590258 | 24614358 | n/a | n/a | n/a | n/a | n/a | 6,7 | 5,9 |
| 12 | 33166145 | 33239783 | tmem178b | Transmembrane protein 178B | membrane | n/a | n/a | 8 | n/a |
| 13 | 13170434 | 13279908 | ptprg | Protein tyrosine phosphatase receptor type G | membrane | carbonate dehydratase activity, zinc ion binding | n/a | 7,11 | n/a |
| 13 | 23851482 | 23861896 | fam219aa | protein FAM219A | n/a | n/a | n/a | 8 | n/a |
| 13 | 24060216 | 24085468 | loxl2a | lysyl oxidase homolog 2A-like | extracellular space, membrane | copper ion binding, protein-lysine 6-oxidase activity, scavenger receptor activity | peptidyl-lysine oxidation, sprouting angiogenesis | 8 | n/a |

| | | | | | | | | | |
|---|---|---|---|---|---|---|---|---|---|
| 13 | 25639751 | 25781707 | cpne5b | copine-8-like | n/a | calcium-dependent phospholipid binding Source | n/a | 1,10 | 1 |
| 13 | 26328980 | 26331331 | opn1lw1 | red-sensitive opsin | membrane | G protein-coupled receptor activity, photoreceptor activity | phototransduction, visual perception | 1,2,4,10,11 | 2,6,7 |
| 15 | 2410998 | 2418594 | n/a | n/a | n/a | n/a | n/a | 7,8 | n/a |
| 15 | 22766186 | 22824294 | n/a | transcription factor COE3-like | nucleus | DNA binding, DNA-binding transcription factor activity, metal ion binding | n/a | n/a | 3 |
| 16 | 17940301 | 17944614 | bcap29 | B cell receptor associated protein 29 | endoplasmic reticulum membrane | endoplasmic reticulum to Golgi vesicle-mediated transport, intracellular protein transport, protein localization to endoplasmic reticulum exit site | n/a | n/a | 10 |
| 17 | 10115782 | 10123352 | n/a | cytosolic sulfotransferase 3-like | n/a | sulfotransferase activity | n/a | 1 | n/a |
| 17 | 12572258 | 12606006 | b4galt5 | UDP-Gal:betaGlcNAc beta 1,4-galactosyltransferase, polypeptide 5 | Golgi cisterna membrane, Golgi membrane | glycosyltransferase activity, metal ion binding | carbohydrate metabolic process, dorsal/ventral axis specification, otolith development, positive regulation of BMP signaling pathway, protein glycosylation, proteoglycan biosynthetic process | 5,10,11 | 11 |
| 20 | 602637 | 626056 | CASKIN2 | caskin-2-like | n/a | n/a | n/a | 1,5,8,10 | 1,6,7,8 |
| 20 | 21206539 | 21254556 | n/a | n/a | n/a | n/a | n/a | 4 | 9 |
| 22 | 3170729 | 3184972 | zbtb34 | zinc finger and BTB domain-containing protein 34-like | n/a | n/a | n/a | 4 | n/a |
| 22 | 3578170 | 3614889 | glt1d1 | Glycosyltransferase 1 domain containing 1 | n/a | glycosyltransferase activity | n/a | 3 | 7,8,10 |
| 22 | 5992293 | 6522529 | grid2 | Glutamate receptor, ionotropic, delta 2 | postsynaptic membrane | ionotropic glutamate receptor activity | n/a | 2 | n/a |
| 22 | 10654447 | 10673831 | serinc5 | Serine incorporator 5 | plasma membrane | n/a | n/a | 7 | 5 |

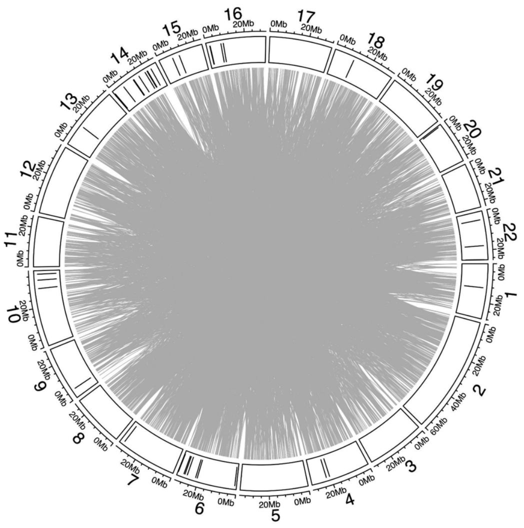


**Fig. S1** Multispecies inter-chromosomal LD versus segregation distortion in the three crosses. The outer tract shows the locations of the 50 kb windows that contained both a segregation distortion SNPs from either of the three crosses and a SNP in high multispecies inter-chromosomal LD ($r^2$>0.5). The gray lines in the middle indicate the connections between the SNPs in multispecies inter-chromosomal LD pairs with an $r^2$>0.5.

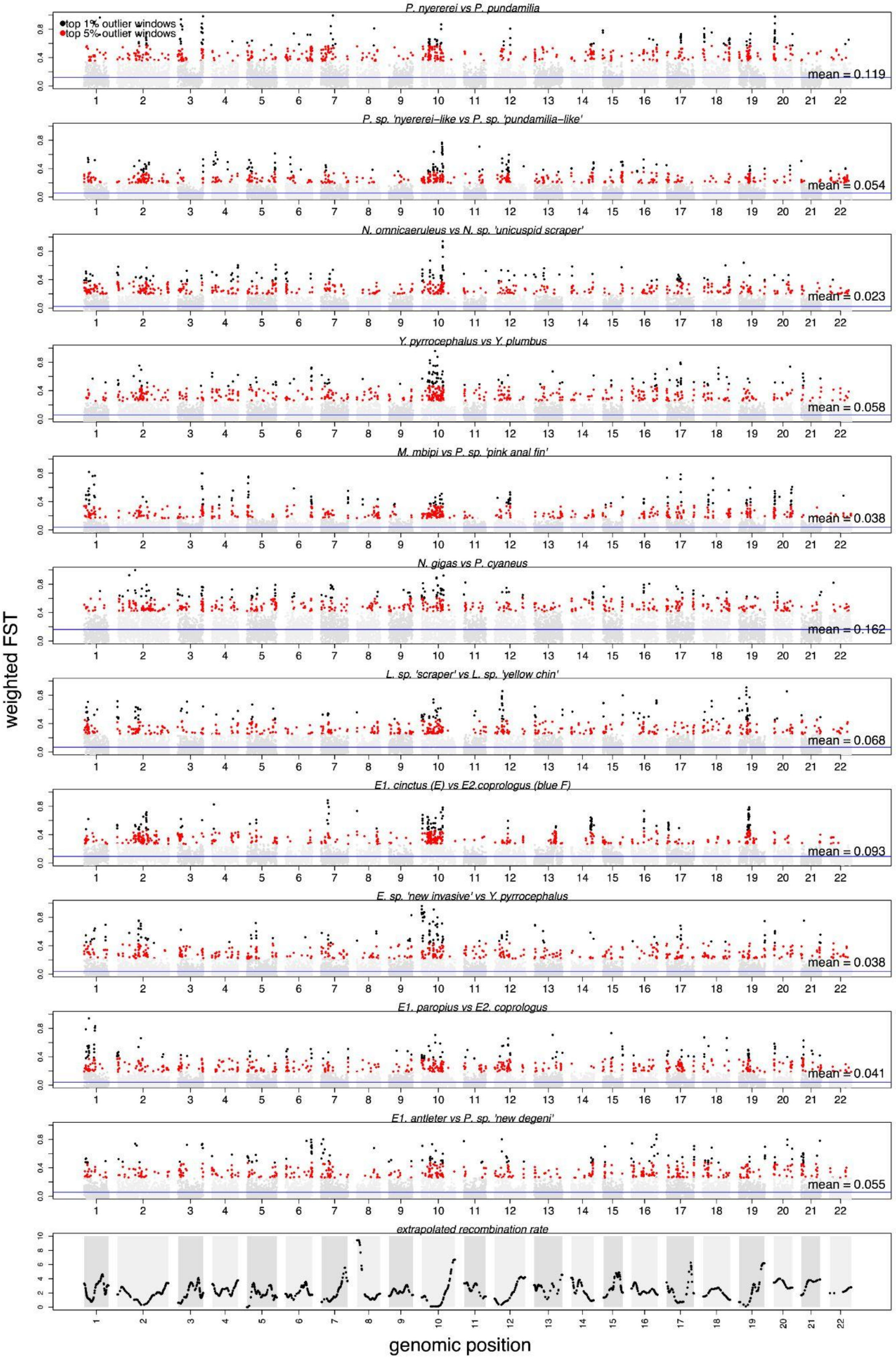


**Fig. S2.** FST landscapes between 11 sympatric sister species pairs (see also Table S4) and recombination landscape inferred from the *P. pundamilia* x *P.* sp. 'red-head' cross by Feulner *et al.* (2018). The FST landscapes are highly correlated with each other as well as with the recombination landscape (Table S7).

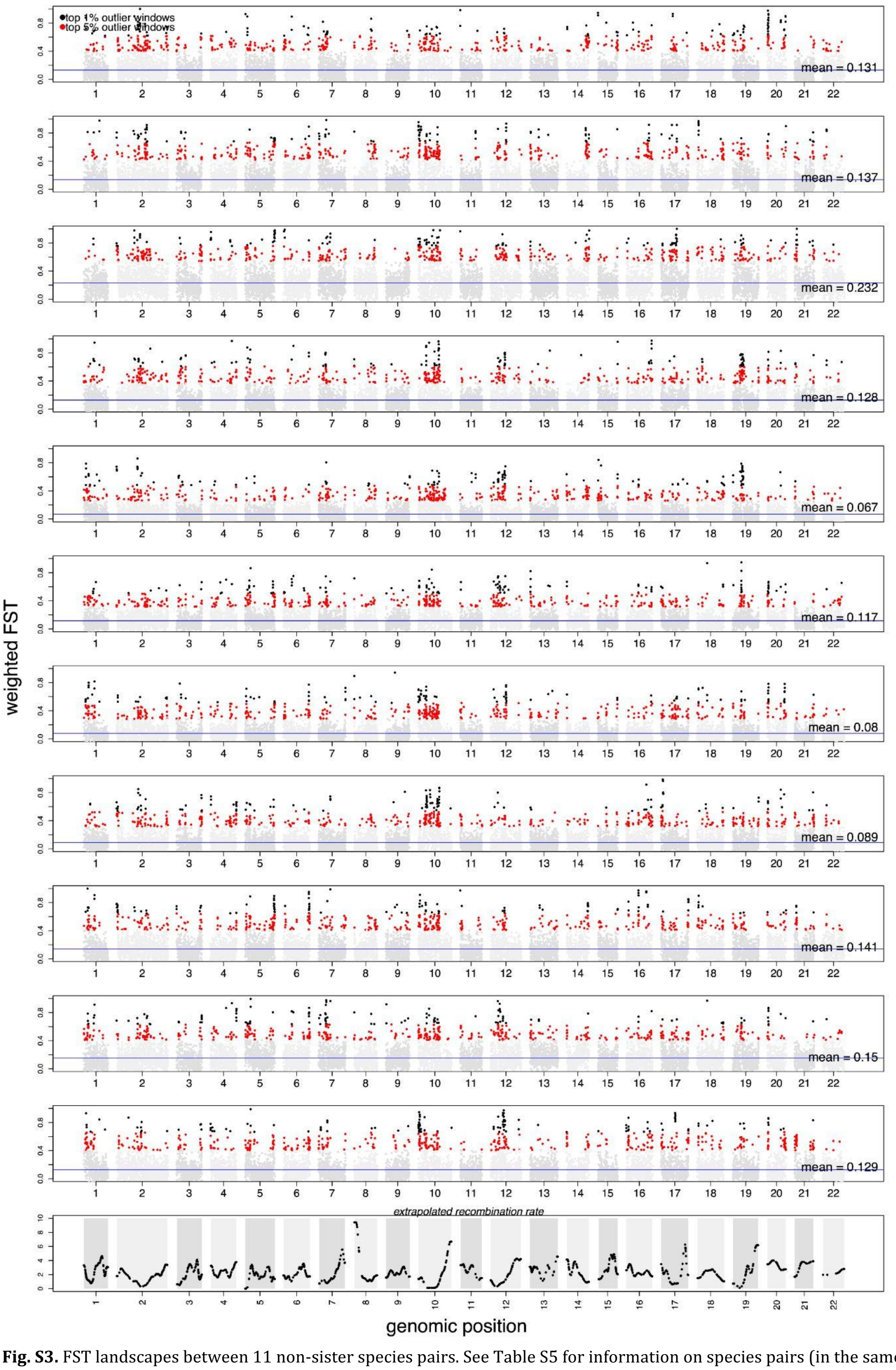


**Fig. S3.** FST landscapes between 11 non-sister species pairs. See Table S5 for information on species pairs (in the same order).
